# Inflammatory Fibroblasts Promote Repair After Injury Through Epithelial Proliferation

**DOI:** 10.64898/2026.08.02.742235

**Authors:** Luis R. Rodríguez, Willy Roque Barboza, Aditi Murthy, Niran Hadad, Sarah Bui, Dakota L. Jones, Yaniv Tomer, Charlotte H. Cooper, Ana Reineberg, Rea Chroneos, Evan T. Hoffman, Surafel Mulugeta, Jeremy Katzen, Jonathan A. Kropski, Nicholas E. Banovich, Michael F. Beers

**Author notes:** M.F.B. is the corresponding author. These authors provided equal contribution to this manuscript (Luis R. Rodríguez, Willy Roque Barboza). **Lead Contact Author:** Michael F. Beers, M.D., Pulmonary and Critical Care Division, Perelman School of Medicine at The University of Pennsylvania, Edward J Stemmler Hall Suite 216, 3450 Hamilton Walk, Philadelphia, Pennsylvania 19104-6118.

## Abstract

Fibroblast heterogeneity after lung injury is a well observed phenomenon made highly relevant by the widespread application of single cell RNA-sequencing. The characterization of homeostatic and injury associated states has led to the identification of a population of fibroblasts that emerge during inflammation and express cytokines that may potentially amplify the inflammatory circuit. However, whether these cells actively contribute to inflammation or serve an alternative function within the broader injury-repair cascade remains unclear. By integrating several robust murine lung injury data sets we establish the persistence of the inflammatory fibroblast across multiple injury models and identify a role for these cells in lung repair after injury through effects on alveolar epithelial proliferation. We validate this observation in-vivo using a genetic model of spontaneous lung fibrosis and in-vitro with mixed alveolar organoid cultures of various homeostatic and injury associated fibroblasts wherein we identify a mesenchymal-epithelial BMP signaling axis as a key driver of the AT2 cell injury repair response. Finally, we present supporting evidence from human disease, reinforcing the relevance of this fibroblast subset in pathological settings. These findings extend critical observations made prior to the single-cell era and contribute to our evolving understanding of fibroblast heterogeneity as a key feature of lung repair.

## INTRODUCTION

Our understanding of mesenchymal biology during lung injury and repair has greatly accelerated with the emergence of single cell technology and the expansion of bioinformatic tools to explore these data sets. Specifically, the characterization of pulmonary fibroblasts has progressed from a simplistic classification of healthy fibroblasts and injury-associated myofibroblasts to a nuanced understanding of specialized subsets of fibroblasts, each with distinct names and biological significance. These studies have led to the generation of integrated atlases from human^1–5^ and mouse^3,5^ data partially aiming to aid the field via the application of an uniform nomenclature. A central theme across these investigations, from the earliest single-cell analyses^6^ to more recent efforts^7–10^, is the pivotal role that specific fibroblast populations play in supporting the pulmonary epithelium during injury. Furthermore, beyond injury and repair, lung fibroblast heterogeneity is highly pertinent to homeostatic and developmental lung biology as the role for fibroblast subpopulations in direct support of epithelial development is clear ^6,8,9,11,12^.

The integration of these scRNA-seq transcriptomics data sets with additional studies incorporating lineage tracing, functional in vitro models, and spatial transcriptomics ^7,8,10,13–15^, has established some consensus in the description of three homeostatic lung fibroblast populations: Alveolar/AF1, Adventitial/AF2, and Peribronchial Fibroblasts. Injury-associated fibroblasts, on the other hand, are less widely agreed upon with varying nomenclature that continues to challenge the field. Much of this can be attributed to the widely used term “myofibroblast” in the description of activated, smooth muscle actin (ASMA) expressing, collagen producing fibroblasts seen throughout the injured lung^16–18^. Adding to this confusion is the highly contractile, ASMA positive fibroblasts present during lung development that are critical to the formation of alveoli but are no longer seen in the adult lung^6,7,10,19^. However, recent clarity has emerged with the description of Collagen Triple Helix Repeating Coil 1 (*Cthrc1)* as a transcriptional marker of pathological fibroblasts in murine models of lung fibrosis^20^. This *Cthrc1*+ population, dubbed the fibrotic fibroblast, was also reported in other murine injury models^21–24^ and human disease^2,14–16,25^ data sets which have been followed by the careful application of lineage tracing to implicate the alveolar fibroblast as the likely cell of origin for this pathological fibrotic fibroblast^26,27^. However, the question of how a supportive fibroblast enters a pathological state and what signaling pathways may promote this disruption are incompletely understood.

A potential avenue to approach this question arose with the description of an injury associated fibroblast population that seems to precede the fibrotic fibroblast called either “transitional” or “inflammatory” fibroblasts ^22,23,26,28^. The name inflammatory refers to the capacity of immune cell derived ligands such as Il-1ß ^26^ or Prostaglandin F2α ^22^ to induce the inflammatory transcriptional program in alveolar and adventitial fibroblasts *in-vitro* as well as the demonstration of enhanced transcriptional expression of a wide range of cytokines in this population ^23,26,29^. Conversely, the name transitional stems from bioinformatic evidence that these fibroblasts occupy an intermediate state between homeostatic and fibrotic fibroblasts^23,28^. Regardless of nomenclature, the biological relevance of this population and its potential to serve as either a proinflammatory cell, a progenitor pool for fibrotic fibroblasts, or some other undefined role is a timely high impact question.

To address this question, we turned to our genetic mouse model of spontaneous lung fibrosis which initiates intrinsic alveolar epithelial stress from expression of disease relevant *SFTPC* mutations to induce acute alveolitis followed by widespread non-resolving fibrotic remodeling ^30^. This model provides a widespread lung injury that thoroughly disrupts pulmonary fibroblast homeostasis, resulting in severe reductions in homeostatic alveolar and adventitial populations with emergence of both inflammatory and fibrotic fibroblast populations ^22,31^. Combining this *in-vivo* model with large scale integration of publicly available human and mouse data sets we interrogated the functional role of the inflammatory fibroblast and found a role for both TGFb and BMP signaling in these events. Further, we use careful reductionist organoid modeling to validate an exciting role for inflammatory fibroblasts as mediators of epithelial proliferation after lung injury. These data support the novel concept for a protective role of a previously underrecognized mesenchymal cell which participates in intricate crosstalk with epithelial cells in the injured and fibrotic niche while providing a conceptual framework for considering new pathways, cells, and mediators as therapeutic targets.

## RESULTS

### Inflammatory Fibroblasts Emerge in the Murine Lung After Injury

We initiated this study by first integrating a wide range of data sets from recent preclinical models characterizing mesenchymal population dynamics during influenza infection ^32,33^ and exogenous bleomycin injury ^20,21,26,27,32^ with data from our *Sftpc* genetic model of spontaneous ^22,31^ lung fibrosis (both published^22,31^ and the current study) as well as naive mice ^34^. Unsurprisingly, after integration most cells expressed mesenchymal markers, though we were able to identify and exclude a minority of cells from the immune, epithelial, and endothelial cell compartments (**Supplemental Figure 1**). UMAP analysis of epithelial, immune, and endothelial lineage negative (*Pecam1-*, *Ptprc-*, *Epcam-*) mesenchymal cells resulted in the identification of 10 unique clusters (**Figure 1A**) all previously described in recent literature ^22,23,26,32^. Though not every population was present across every study, we did not identify any population unique to a specific publication, supporting the validity of the integration and the quality of the data (**Figure 1B; Supplemental Figure 2**). This integration allowed us to further validate markers that have been consistently reported across the literature ^6,20,22,23,26,27,32,34,35^ for all the mesenchymal states, including the stress activated state ^20,26^ that up to this point has only been reported in bleomycin injury (**Figure 1C; Supplemental Figure 2**). Notably, we now report the emergence of the “stress activated” state after flu injury (**Figure 1D**). Furthermore, this integrated analysis demonstrates the limited presence of fibrotic fibroblasts in non-fibrotic injury models and supports the application of “fibrotic” nomenclature to this population (**Figure 1D**). Importantly, we note that a robust number of inflammatory fibroblasts emerge across all injury models and, as was recently noted for the fibrotic fibroblast ^26,27^, appear to be derived almost entirely from *Pdgfra+* alveolar fibroblasts expressing markers *Scube2* and *Lepr* (**Figure 1E**). These findings highlight a conserved response to lung injury in mice, where a shared signaling niche promotes the emergence of inflammatory fibroblasts across both resolving and non-resolving lung injuries.

**Figure 1.**
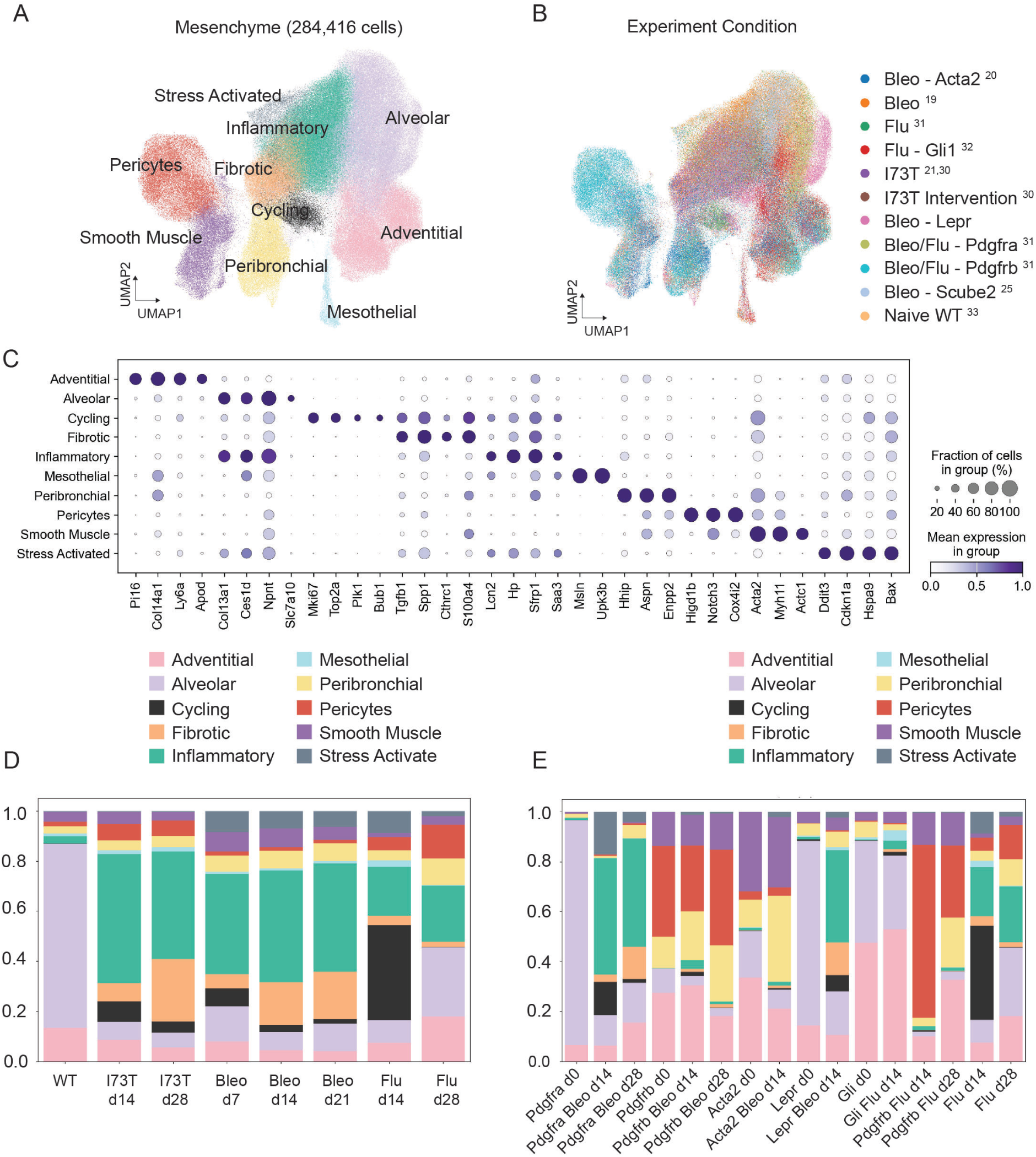
Inflammatory Fibroblasts Appear Across Multiple Lung Injuries: A) UMAP of integrated data sets (GSE253453, GSE221402, GSE132771, GSE234604, GSE296513, GSE215094, GSE248798, GSE244237, GSE249931, GSE229523, GSE276546, GSM6428697, as well as the data generated in this study) including 248,416 mesenchymal cells from flu, bleomycin, *Sftpc*^I73T^ lung injuries and naïve mice clustered and annotated. B) UMAP representation of the same 284,416 cells classified according to injury type and lineage tracing approach. Subscript in the legend indicates reference citation. C) Gradient dot plot of marker genes used to identify the annotated clusters. D) Stacked bar plots of the frequency from each cluster during specified time points in each injury model. E) Stacked bar plots of the cluster frequency from each lineage trace experiment clustered during specified time points in the injury models.

### TGFß Inhibition Inhibits Fibrotic Remodeling, Preserves Inflammatory Fibroblast Frequencies, and Supports Epithelial Expansion

We and others have previously demonstrated that endogenous TGF-ß1 is capable of driving expression of the fibrotic fibroblast transcriptional program in cultured fibroblasts ^20,22,23^. Additionally, recent work using an alveolar fibroblast specific Cre recombinase to inhibit TGF-ß signaling in alveolar fibroblasts during bleomycin injury has demonstrated that TGF-ß is necessary for the emergence of fibrotic fibroblasts, but not inflammatory fibroblasts ^26^. These observations indicate that TGF-ß is a critical driver of the inflammatory/fibrotic switch. Thus, to explore the effects of inflammatory fibroblast persistence after lung injury we turned to our *Sftpc*^I73T^ genetic model of lung fibrosis and intervened 12 days after tamoxifen induction using the ALK5 inhibitor (ALK5i), SB525334, to block TGF-ß1 receptor signaling (**Figure 2A**). ALK5i protected mice from fibrotic injury with significantly reduced weight loss and mortality along with normalization of static lung compliance (**Figure 2B-C**). This was accompanied by decreased bronchoalveolar lavage fluid (BALF) markers of lung injury and normalization of soluble collagen content (**Figure 2I-G**). Assessment of lung histology also demonstrated a quantitative decrease in lung injury and fibular collagen deposition after ALK5i intervention (**Figure 2H-I**). Gene expression analysis of whole lungs lysate 28 days post tamoxifen confirmed the expected decrease in the fibrotic fibroblast signature as well an increase in alveolar, adventitial, and inflammatory fibroblast signatures after ALK5i (**Figure 2J**).

**Figure 2.**
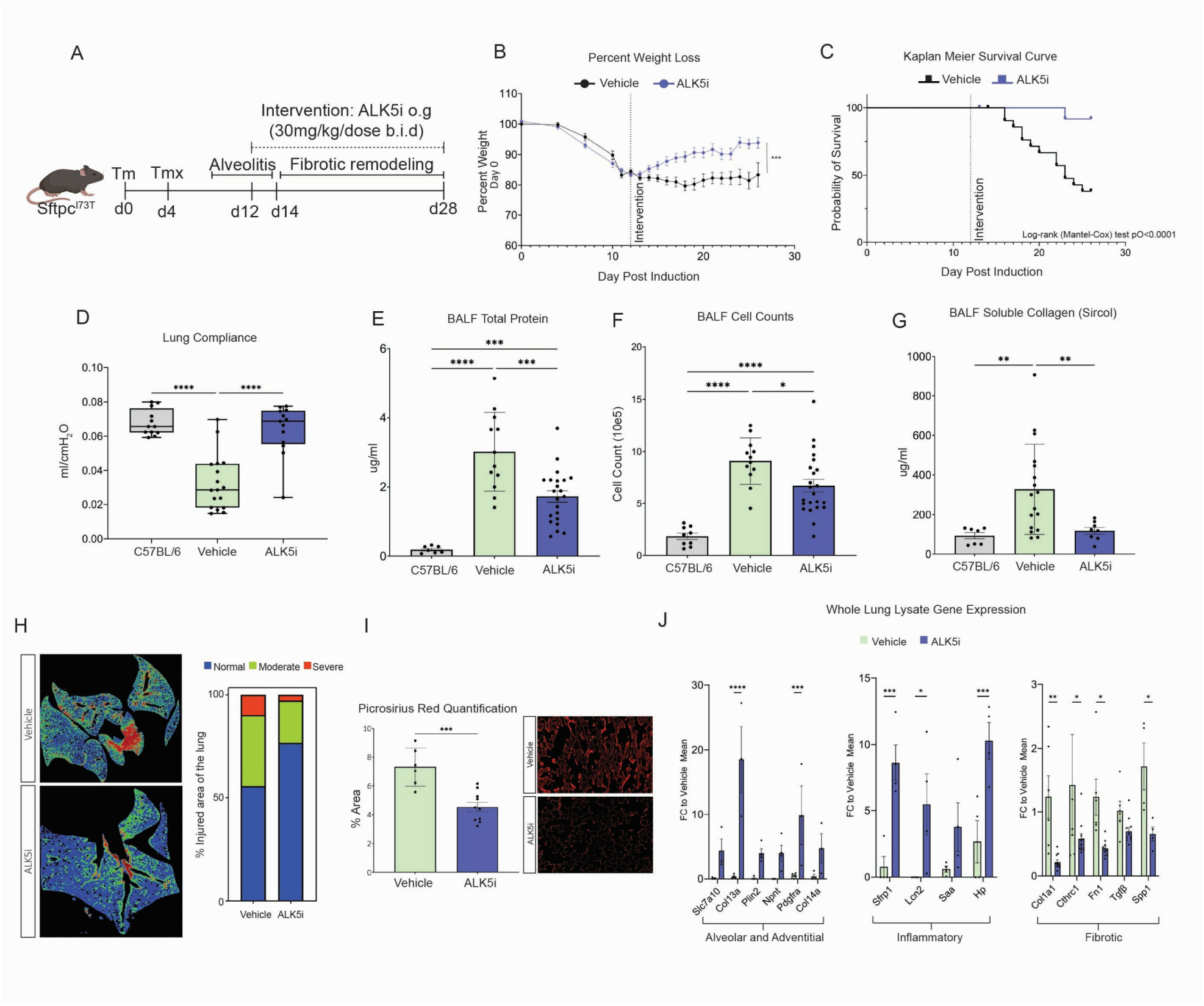
TGF-β1 Receptor Inhibition Resolves Fibrotic Injury: A) Schematic of intervention protocol outlining tamoxifen delivery to induce expression of SPC-C^I73T^ at days 0 and 4 followed by daily intervention with oral gavage (o.g.) ALK5 inhibitor SB525334 (60 mg/kg BID, n = 20 mice per group). B) Weight loss was significantly decreased after Alk5i intervention. C) 28-day mortality was significantly reduced in ALK5i group. D) Static lung compliance, measured by Scirec Flexivent, increased significantly in ALK5i group compared to vehicle control (mean ± SEM; n=13 for vehicle and ALK5i group and n=12 for WT group. E-G) Markers of lung injury, inflammation, and fibular collagen in the bronchoalveolar lavage (BAL) fluid collected from mice at 28d post tamoxifen were significantly reduced in ALK5i-treated mice (mean±SEM; n=12 mice per group). H) Representative histology with fibrosis severity scoring algorithm (normal = blue, moderate = green, and severe = red) and bar graph showing relative amounts of injury per lung lobe demonstrating a significant p = 0.0162 increase in normal lung after ALK5i (n= 9 per group). I) Picrosirius staining of fixed lung sections demonstrates significant decrease in fibrillar collagen deposition after ALK5i (mean ± SEM; n=10 for ALK5i n=6 for vehicle). J) Gene expression analysis via QPCR in whole lung lysates of marker genes associated with alveolar, adventitial, inflammatory, and fibrotic fibroblasts (mean ± SEM; n=4 per group). *p<0.05, **p<0.005, ***p<0.0005, ****p<0.00005 by ordinary one-way ANOVA.

Having seen this pronounced effect on gene expression associated with lung injury and aberrant repair, we elected to perform scRNA-seq after ALK5i to evaluate fibroblast and epithelial population dynamics. To align with the physiology and fibrotic data collected (**Figure 2**) we chose the 28-day time point after tamoxifen and 16 days post starting ALK5i for the sequencing analyses (**Figure 2A**). Given the focus on epithelial and mesenchymal populations, our single cell digestions of mouse lungs were first CD31 and CD45 depleted. Following this we bead sorted CD326+ epithelial cells and performed a single cell analysis on a cell suspension comprised of 25% epithelium (CD326+) and 75% trilineage negative (CD31-, CD45-, CD326-) mesenchyme (**Supplemental Figure 3A**). Using previously defined markers for pulmonary epithelial and mesenchymal subsets (**Supplemental Figure 3B-D**) we identified changes to fibroblasts and alveolar epithelial cell frequencies after ALK5i (**Figure 3A-B**). Fibrotic fibroblast frequency decreased, and alveolar fibroblast frequency increased in the ALK5i samples. Interestingly, there was little change in the frequency of inflammatory fibroblasts while the epithelial data presented an increase in activated AT2 cells. Further, as with the fibrotic endpoints from Figure 2, we also observed a transcriptomic decrease in fibrosis associated genes (**Figure 3C**).

**Figure 3.**
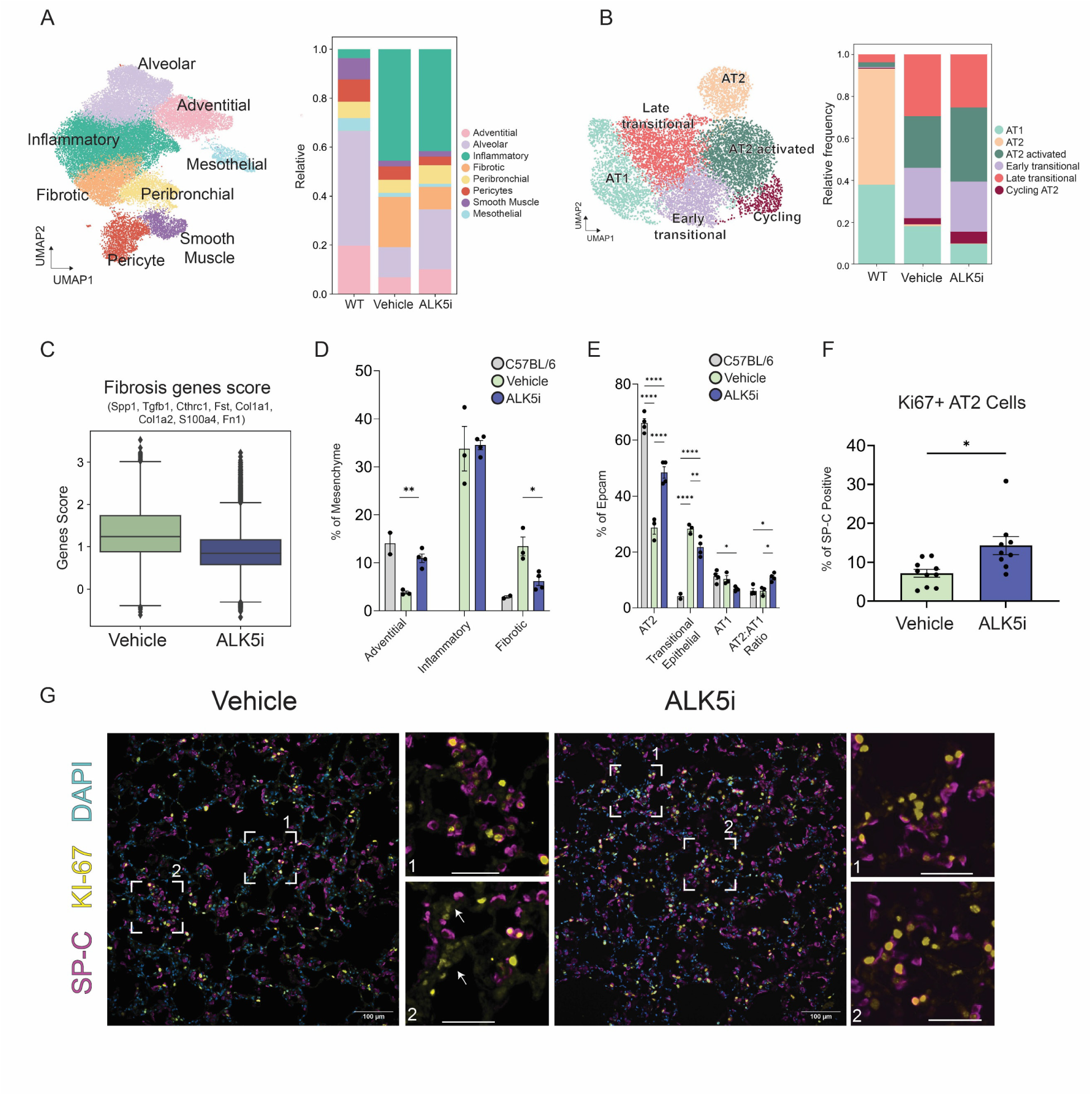
AT2 Proliferation Increases After TGF-β2 Receptor Inhibition: A) UMAP visualization of 35,695 mesenchymal cells color-coded by cell lineage along with frequency of each cluster in each of the integrated conditions (n = 2 mouse lungs per condition) B) UMAP visualization of 10,117 distal alveolar cells color-coded by cell lineage along with frequency of each cluster in each of the integrated conditions (n = 2 mouse lungs per condition). C) Fibrosis gene score is significantly decreased after ALK5i. D-E) Flow cytometry quantification of whole lung single cell suspension identifies an decrease in injury associated fibrotic fibroblasts and transitional epithelial cells while also showing and increase in homeostatic adventitial fibroblasts and AT2 cells in the ALK5i group relative to vehicle (mean±SEM; n=3-4 mice per group). F) Lung sections were stained for Ki67 and SP-C expression followed by quantification of double positive cells as a percentage of total SP-C positive cells. Quantification was performed on 5 fields per mouse (n = 3 mice per condition). G) Representative immunofluorescent staining of lung sections from vehicle or ALK5i treated *Sftpc*^I73T^ 28 days post tamoxifen induction stained with antibodies against HA, Ki67, and DAPI for nuclei counterstaining. For clarity, as *Sftpc*^I73T^ mice express HA tagged mutant SP-C the images are labelled SP-C rather than HA. Scale bars:100µm. *p<0.05, **p<0.005, ***p<0.0005, ****p<0.00005 by ordinary one-way ANOVA.

The changes in scRNA-seq frequencies, validated using flow cytometry (**Figures 3D-E**), suggested a potential increase in AT2 proliferation under ALK5i. To confirm this, we performed and quantified immunofluorescence co-staining of Ki-67 and SP-C in histological sections from *Sftpc*^I73T^ lungs at 28 days post tamoxifen induction (**Figure 3F-G**). These data inform on the potent role for TGF-ß1 signaling in the emergence of fibrotic fibroblasts, confirm, using a different model of injury-repair, a previous report that inhibition of TGF-ß1 signaling does not reduce inflammatory fibroblasts ^26^, and reveal increased AT2 proliferation under these conditions.

### TGF-ß Inhibition Does Not Promote Increased Inflammation During Fibrogenesis

TGF-ß is an essential regulator of lung injury^36,37^ and its effects as an inhibitor of inflammation^38–40^ are also well described. Recent work in bleomycin injury has demonstrated the efficacy of TGF-ß delivery on the inhibition of early inflammatory events^41^. Coupled to this, alveolar fibroblast specific TGF-ß inhibition lead to inhibition of fibrosis but increased alveolar permeability and inflammation^26^. Therefore, to check if our ALK5i was masking the proinflammatory capacity of the inflammatory fibroblast we performed a 6-week study in *Sftpc*^I73T^ mice where ALK5i was administered starting at day 12 post tamoxifen and either removed at day 28 or continued out to day 42 (**Supplemental Figure 4A**). Removal of ALK5i did not result in significant increases in mortality or alterations in static lung compliance (**Supplemental Figure 4B-C**). Total BALF cell counts were not impacted by the removal of ALK5i while interstitial macrophages and T-regulatory cells, key immune populations associated with fibrotic injury, did not increase after ALK5i removal (**Supplemental Figure 4D-F**). These data do not support that inflammatory fibroblasts are a significant contributor to proinflammatory signaling.

### Inflammatory Fibroblasts Support Reparative Alveolar Organoid Formation

While it appeared that inflammatory fibroblasts did not likely contribute to inflammation in this model, we noted that flow cytometry analysis of our extended ALK5i study revealed a continued increase in the number of AT2 cells as well as a decrease in transitional AT2s (**Supplemental Figure 4G-H**). We then hypothesized that the inflammatory fibroblast population may instead participate in the epithelial proliferation observed in vivo (**Figure 3F, 3G)**. To explore this potential interaction, we chose to apply mixed cell alveolar organoid modeling. Protocols to isolate alveolar, adventitial, and fibrotic fibroblasts have been recently described ^20,22,42^, however the isolation of inflammatory fibroblasts has not. In previous scRNA-seq analysis we noted that the alveolar fibroblast population is nearly absent from the *Sftpc*^I73T^ mouse after tamoxifen induction, and we capitalized on this observation to FACS isolate inflammatory fibroblasts (**Figure 4A**). Modifying the published strategies, we then isolated alveolar, adventitial, inflammatory, and fibrotic fibroblasts and analyzed their transcriptomes via population RNA-seq. The resulting populations clustered separately in Principal Component Analysis (**Figure 4B**) and expressed population specific markers confirming that efficacy of the strategies (**Figure 4C**). Pathway enrichment analysis comparing injury associated (fibrotic and inflammatory) fibroblasts to homeostatic (alveolar and adventitial) fibroblasts identifies a signature consistent with increased ECM production and matrix deposition after injury (**Figure 4D, 4E**). Homeostatic populations are enriched in signal transduction and immune system signaling consistent with alveolar and adventitial signaling to epithelial and immune cells respectively (**Figure 4F,4G**). Interestingly, fibrotic fibroblasts have enriched phosphorylated SMAD signaling supporting activation of TGF-ß signaling in fibrotic fibroblasts.

**Figure 4.**
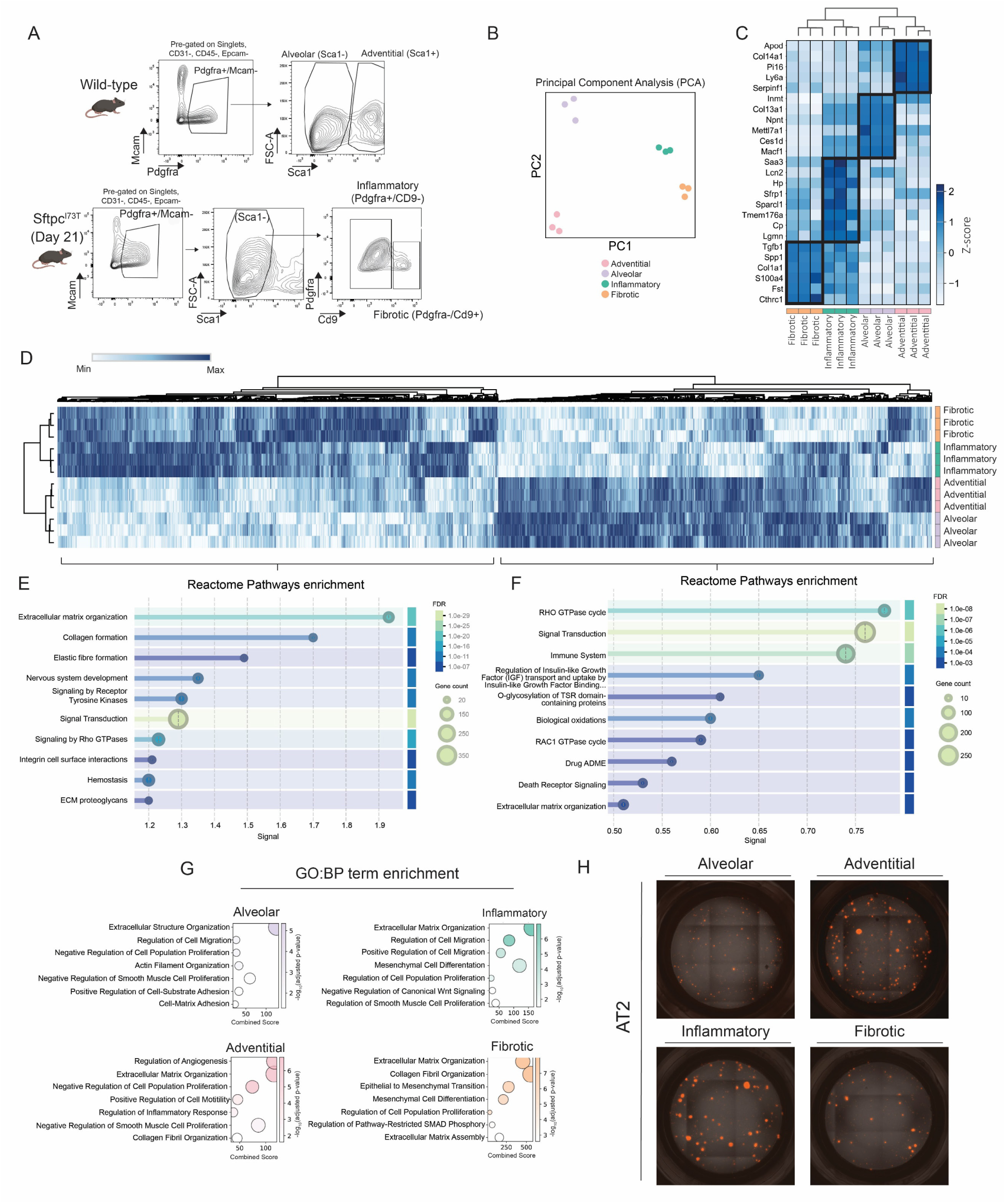
Characterization of Flow Sorted Fibroblast Populations: A) Flow sorting strategy for the isolation of adventitial, alveolar, inflammatory, and fibrotic fibroblasts. B) Principal component analysis of RNA isolated from each of the four fibroblast populations (n=3). C) Unsupervised hierarchical clustering (Euclidean) heatmap (row normalized z-score) of select marker genes for each fibroblast population. D) Unsupervised hierarchical clustering (Euclidean) heatmap (row normalized z-score) of all differentially expressed genes (FDR <0.05) E) String DB summary Reactome analysis of top 10 terms enriched in fibrotic and inflammatory fibroblasts. F) String DB summary Reactome analysis of top 10 terms enriched in alveolar and adventitial fibroblasts. G) Gradient dot plots of top 7 gene ontology (GO) terms from biological process (BP) enriched in differentially expressed genes from each fibroblast population. The size of circles reflects the magnitude of the combined term score. H) Representative fluorescent microscopy images of 14d organoid cultures derived using Tdt^+^ AT2 cells from *Abca3*^Cre^*Rosa26*^Tdt^ mice and the four fibroblasts population isolated from WT or *Sftpc*^I73T^ mice at 21d post tamoxifen induction. Scale bars:500µm.

Once the isolation strategy was established, we were next able to functionally characterize the various fibroblast subsets. In agreement with the literature ^6,34,43^, we observed increased organoid area and colony numbers in homeostatic co-cultures of adventitial fibroblasts as compared to alveolar fibroblasts (**Figure 4H, Supplemental S5A**). Interestingly we observed that inflammatory fibroblasts supported the formation of alveolar organoids to a greater degree than alveolar or fibrotic fibroblasts while fibrotic fibroblasts did not (**Figure 4H, Supplemental S5A**). We performed these same experiments with activated AT2s derived from our fibrotic model and, despite the decreased progenitor capacity of activated AT2s ^44^, we again observed that inflammatory fibroblasts supported organoid formation rescuing the defective AT2 phenotype (**Supplemental S5A**). These observations along with the results of our *in-vivo* ALK5i studies indicate that inflammatory fibroblasts induce AT2 proliferation during lung injury.

### Inflammatory Fibroblasts Promote AT2 Proliferation via BMP Inhibition

Informatic analysis of scRNAseq dataset for cell-cell cross talk indicated that inflammatory fibroblasts were producing ligands that directly interacted with activated AT2 cells, particularly after ALK5i (**Figure 5A**). Thus, to define the potential signaling mechanism driving inflammatory fibroblast mediated AT2 proliferation we interrogated our population RNA-seq and sc-RNA-seq data for ligands that have previously been reported to drive AT2 expansion during injury. One major pathway that emerged and was of significant interest was BMP signaling. This pathway is comprised of multiple TGF-ß superfamily members, has abundant previous literature suggesting its relevance in the lung^45–49^, and its crosstalk is altered in our mouse injury after ALK5i highlighted by the decrease in BMP signaling between alveolar and activated AT2 cells(**Figure 5B**). This transcriptomic analysis identified several BMP antagonists upregulated in both inflammatory and fibrotic fibroblasts while alveolar fibroblasts expressed high levels of BMP4 (**Supplemental Figure 5B**). Furthermore, we also noted that TGF-ß1 and -ß2 were both upregulated in fibrotic fibroblasts (**Supplemental Figure 5C**). We previously demonstrated that the addition of TGF-ß1 to inflammatory fibroblasts promoted the entry into the fibrotic state^22^. In our organoid cultures exogenous TGF-ß1 was sufficient to inhibit alveolosphere formation and expansion (**Supplemental Figure 5D**), suggesting that TGF-B in the injury milieu can override any proliferative signaling.

**Figure 5.**
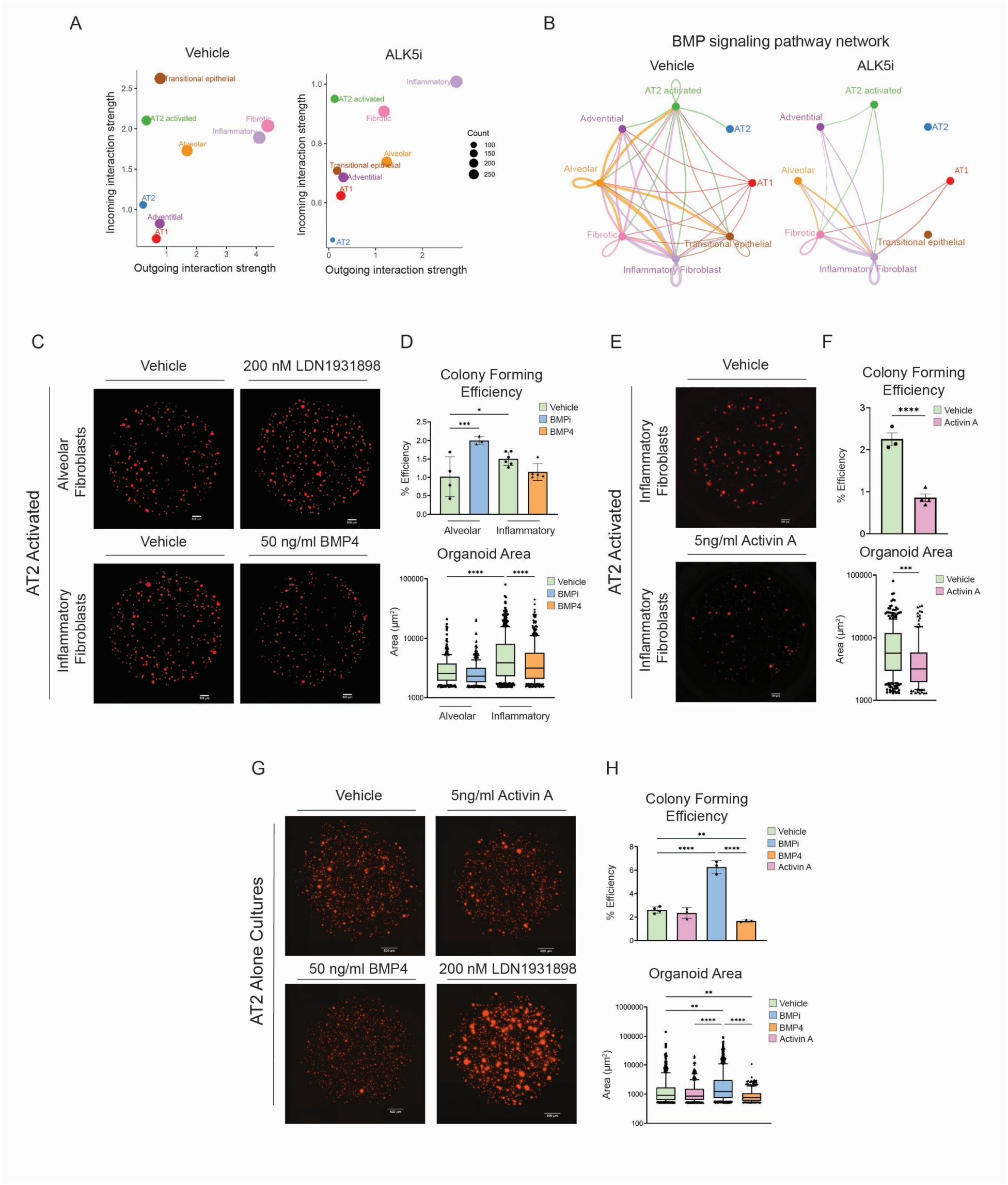
Inflammatory Fibroblast Derived BMP Antagonists Promote AT2 Proliferation: A) Cell Chat visualization of interactions between distal epithelial and mesenchymal clusters belonging to the Cell Chat BMP signaling pathway term. B) Dot plot visualization of receptor ligand interactions between distal epithelial and mesenchymal clusters with x and y axis denoting outgoing and incoming interaction strength respectively. C-F) Representative fluorescent microscopy images of 14d organoid cultures derived using Tdt^+^ activated AT2 cells from *Abca3*^Cre^*Rosa26*^Tdt^*Sftpc*^I73T^ mice and either alveolar or inflammatory fibroblasts isolated from WT and *Sftpc*^I73T^ mice respectively at 14d post tamoxifen induction. Organoid diameter (μm) and colony forming efficiency (CFE; %) of organoids. Organoids were treated with BMP inhibitor LDN1931898 (200nM), BMP4 (50ng/ml), Activin A (5ng/ml), or vehicle control refreshed every 48 hours. G, H) Representative fluorescent microscopy images of AT2 cell monoculture organoids derived from *Abca3*^Cre^*Rosa26*^Tdt^*Sftpc*^WT^ mice and cultured for 14 days. Quantification and challenge conditions are the same as above. I) Graphical summary of signaling between fibroblast populations and AT2 cells leading to AT2 proliferation during injury repair. Scale bars:500µm. *p<0.05, **p<0.005, ***p<0.0005, ****p<0.00005 by ordinary one-way ANOVA.

Given the previous literature reporting BMP4 as an inhibitor of AT2 proliferation ^45^ we turned back to organoid modeling to validate fibroblast specific regulation of AT2 proliferation in the post-natal lung by manipulation of BMP signals. To align with our *in-vivo* studies using a lung injury model, our organoids were performed using activated AT2s derived from the *Sftpc*^I73T^ mouse 14 days after tamoxifen induction. We first exposed BMP4 expressing alveolar fibroblasts to the BMP inhibitor (BMPi) LDN1931898 and BMP4 antagonist expressing inflammatory fibroblasts to exogenous BMP4. Under these conditions BMPi promoted proliferation of AT2 cells despite the presence of alveolar fibroblasts and, in turn, exogenous BMP4 inhibited the enhanced organoid formation associated with the inflammatory fibroblast co-culture (**Figure 5C; 5D**). Exogenous BMP4 has been shown to inhibit AT2 proliferation ^45^, but our hypothesis was that BMP antagonists released by inflammatory fibroblasts are inhibiting tonic alveolar fibroblast derived BMP4 present during homeostasis. To directly test this, we added a low concentration of activin A to inflammatory fibroblasts co-cultures which binds BMP antagonists thereby inhibiting their activity. The addition of activin A attenuated the formation of alveolar organoids in the presence of inflammatory fibroblasts (**Figure 5E-F**). Finally, the addition of low level activin A to AT2 monocultures does not inhibit alveolosphere formation, confirming that at this concentration, activin A does not directly act on AT2 cells to inhibit proliferation (**Figure 5G-H**). Collectively, these data suggest that under homeostasis alveolar fibroblasts constitutively release BMP4 to inhibit AT2 proliferation, however upon injury, inflammatory fibroblasts emerge and begin to secrete BMP4 inhibitors, suppressing the local BMP4 inhibition signal, “releasing the brakes”, and allowing for AT2 proliferation.

### Human Inflammatory Fibroblast Transcriptome Suggests a Role for Proliferation During Repair

Having utilized preclinical mouse models to establish a function for inflammatory fibroblasts in repair after lung injury via BMP signal modulation, we asked if there was a similar role for these cells in human disease. Inflammatory fibroblasts have been described in both IPF^14,23,26,50^ and Hermansky-Pudlak Syndrome^29^ but, as with the mouse data, their function is not well described. We again capitalized on publicly available data and interrogated the Human Lung Cell Atlas (HCLA) which consists of over 2 million pulmonary cells across 49 different data sets^1^. Specifically we analyzed the mesenchyme (**Supplemental Figure 6**) and re-annotated based on marker genes that have been used to try and unify the nomenclature of pulmonary fibroblasts across humans and mice^7,14,23,26,50^. We annotated 7 unique clusters of mesenchyme that are found across 7 different lung pathologies with specific transcriptomic signatures (**Figure 6A-C**). As with the mouse data, fibrotic fibroblasts are present in high relative frequencies across fibrotic diseases and much lower frequencies in primarily inflammatory diseases (**Figure 6D**). Of particular note is the substantive presence of fibrotic fibroblasts in COVID-19 lungs, an observation consistent with the risk of secondary fibrosis developing in severe cases^51,52^. Using the top 50 marker genes for the adventitial and alveolar clusters we see that inflammatory fibroblasts have low level expression of both signatures (**Figure 6E**). While we did not identify a similar pattern of BMP4 and BMP antagonists in human alveolar and inflammatory fibroblasts respectively (**Supplemental Figure 6**), inflammatory fibroblasts and adventitial fibroblasts are enriched in transcripts that are associated with AT2 proliferation such as FGF2 and FGF7 (**Figure 6F; Supplemental Figure 6**). Finally, paralleling the mouse inflammatory fibroblast transcriptomic data reported in literature^26^, gene set enrichment analysis confirms that human inflammatory fibroblasts are enriched for immune signaling transcripts and cytokines. Together this analysis suggests that human inflammatory fibroblasts are a pan-injury associated population with the potential to promote epithelial cell proliferation.

**Figure 6.**
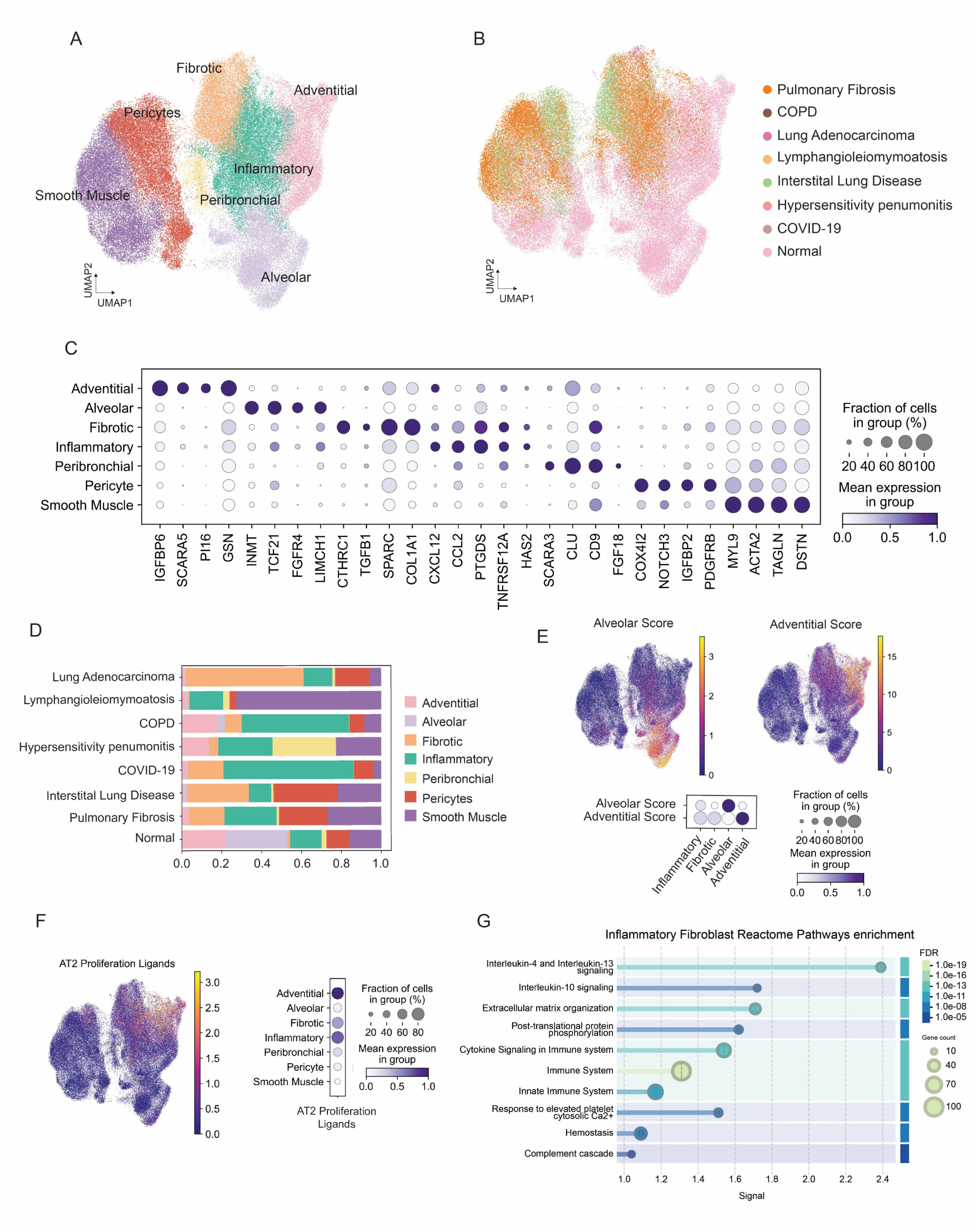
Human Inflammatory Fibroblasts May Support AT2 Proliferation: A) UMAP visualization of human mesenchymal cells from the Human Lung Cell Atlas and color-coded by cell lineage B) UMAP visualization of the same mesenchymal cells color-coded by disease status. C) Gradient dot plot of marker genes used to identify each mesenchyme cluster. D) Stacked bar plots of mesenchyme cluster frequency in each disease state E) UMAP representation and gradient dot plot of adventitial and alveolar gene scores derived from the top 50 marker genes of the respective clusters. F) UMAP representation and gradient dot plot of AT2 proliferation ligand gene score. G) Summary Reactome analysis of top 10 terms enriched in inflammatory fibroblast cluster.

### The Fibrotic Niche in End Stage Lung Disease Contains Inflammatory Fibroblasts

Given the hypothetical role for inflammatory fibroblasts in AT2 proliferation we turned to an analysis of spatial transcriptomic data derived from patients with pulmonary fibrosis^14^. As was previously defined, the samples were derived from either unaffected control patients or “less” and “more” affected fibrotic lungs defined by their relative degree of overall pathology^14^ (**Figure 7A**). The original analysis of this data^14^ identified a population of inflammatory fibroblasts, here we expended this population by including all cells belong to the mesenchymal lineage marked by the expression of PTGDS, CCL2, and HAS2 (**Figure 7B**) which reflects the same markers used to identify inflammatory fibroblasts in the HCLA data set (**Figure 6C**). This dataset was previously annotated using transcriptomic cluster analysis to define cell types and identify cellular niches. Inflammatory fibroblasts were present at varying frequencies across all niches, with the highest frequencies in niches C3, C4, and C10 (**Figure 7**). C3, the transitional niche, showed epithelial detachment, contained most KRT5−/KRT17+ cells, and was enriched in fibrotic foci with fibrotic fibroblasts. C4, the fibrotic niche, consisted of moderate to severe fibrosis near lymphatic vessels and was enriched in plasma cells and diverse fibroblast populations, including inflammatory fibroblasts. Notably, within individual niches, C10 had a higher proportion of inflammatory fibroblasts than any other fibroblast subtype. C10, the mesenchymal niche, is located near the subpleural space and is enriched in mesenchymal cells including mesothelial cells, adventitial fibroblasts, and inflammatory fibroblasts.

**Figure 7.**
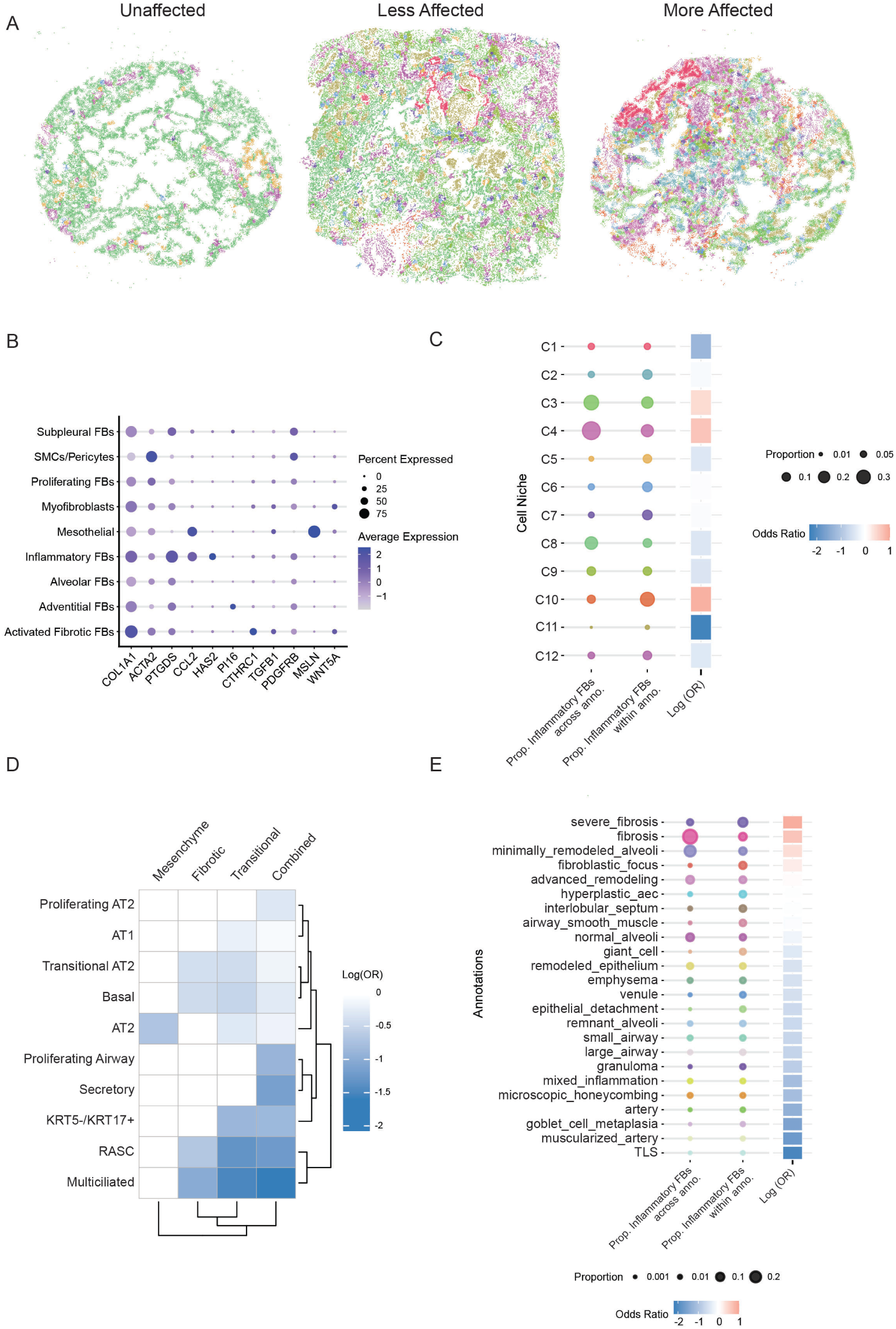
Inflammatory Fibroblasts Are Localized to Lung Injury in Pulmonary Fibrosis: A) Representative examples from unaffected and less/more affected PF samples showing cell-based niches with each point representing a cell centroid. B) Gradient dot plot of marker genes used to identify mesenchyme cell types. C) Proportion of inflammatory fibroblasts distributed across niches (left) and as a proportion of all fibroblasts within a given niche (right). Log OR used to determine the niches with high representation of inflammatory fibroblasts (OR>0 and adjusted p-value < 0.05). D) Heatmap of log odds ratios showing the proximity enrichment of inflammatory fibroblasts with epithelial cell subtypes across niches C3, C4, C10, and all samples combined, with non-significant interactions set to zero. E) Proportion of inflammatory fibroblasts distributed across annotations (left) and as a proportion of all fibroblasts within a given annotation (right). Log OR used to determine the niches with high representation of inflammatory fibroblasts (OR>0 and adjusted p-value < 0.05).

Considering potential epithelial crosstalk, we performed proximity analyses to calculate the odds ratio (OR) between inflammatory fibroblasts and all annotated epithelial populations within the enriched C3 (transitional), C4 (fibrotic), and C10(mesenchymal) niches (**Figure 7D**). Interestingly we observed that, despite the enrichment in pathological niches, inflammatory fibroblasts were not likely to be found proximal to airway cells and the pathological KRT5-/KRT17+ cell. Further analysis across all niches reinforces that inflammatory fibroblasts are most often found in areas that are annotated as fibrosis or some stage of alveoli undergoing remodeling (**Figure 7E**). Together these findings spatially localize the inflammatory fibroblast to areas of potential lung injury and repair.

## DISCUSSION

Our primary goal was to understand the biological function of inflammatory fibroblasts in lung repair after injury and more specifically, define the crosstalk between injury associated fibroblasts and epithelial cells. This is a critical question as studies applying scRNA-seq to various pulmonary injury models and diseases continue to observe complex fibroblast heterogeneity. A secondary goal was to illustrate the significant overlap between data sets across multiple diseases, and to highlight the need to contextualize new data with publicly available data to establish consensus and appropriate biological relevance for fibroblast subsets.

Firstly, the nomenclature of novel cell types in the single cell era requires careful consideration and debate, as is clearly true with the inflammatory fibroblast. In the mouse, this cell state has been defined by markers including *Lcn*2, *Saa3*, *Cxcl12*, and *Sfrp1*^23,26,29^; and it is the expression of cytokines that most clearly connects the human and mouse transcriptomes^26^ (**Supplemental Figure 6**). This is consistent with literature supporting fibroblast to immune cell crosstalk in lung diseases^4,53–60^. Though a recent report suggests that these inflammatory fibroblast cytokines act primarily to amplify inflammation^26^, we did not find an association between immune cell numbers and inflammatory fibroblast frequency (**Figure 3 and Supplemental Figure 3**). Instead, we observed a potential regenerative capacity for inflammatory fibroblasts as drivers of epithelial proliferation *in-vivo* (**Figure 3**) and *in-vitro* (**Figure 4**). Though the mechanistic link we identified did not involve a cytokine (**Figure 5**), there is evidence for cytokines as drivers of epithelial proliferation^6,61–64^. Thus, our data supports the use of the “inflammatory” nomenclature despite the lack of direct evidence for a proinflammatory phenotype in this model.

Our findings also align well with previous observations reporting BMP4 as an inhibitor of epithelial proliferation in murine lungs ^6,45,65,66^ and as a regulator of epithelial biology in lung development^46,67^. However, by identifying the population of fibroblasts that specifically antagonizes this signaling pathway during injury, we advance the understanding of the role of fibroblast heterogeneity during repair. A recent report using *Gli1*^Cre-ERT2^ mice challenged with flu injury suggested that a subset of activated fibroblasts termed “alveolar myofibroblast like” emerged during injury and promoted AT2 expansion^33^. While our analysis of their single cell data suggests that *Gli1* positive adventitial fibroblasts expand and are enriched by their lineage trace (**Figure 1E**), it is possible that their isolation strategy was enriching for inflammatory fibroblasts as well. This further highlights the importance of contextualizing individual single cell studies with integration of publicly available data. Future studies exploring the potential of adventitial fibroblast migration into the alveolar space are necessary within this context.

We were surprised to find that human inflammatory fibroblasts did not express a BMP inhibitory signature as observed in the mouse. It is important to first note that mouse models of lung injury may reflect an early response that is lost or in chronic disease and therefore difficult to observe in advanced disease states such as the end stage IPF lung. However, the consistent expression of ligands associated with epithelial proliferation in human adventitial and inflammatory fibroblasts supports the role of mesenchymal cells as drivers of proliferation during injury. Additionally, there is consistent expression of inflammatory cytokines in the human inflammatory fibroblast population. We reiterate that inflammatory cytokines, such as Il-1b^68^ and Il-6^6^, have been implicated as drivers of epithelial proliferation, suggesting that epithelial proliferation during injury is a coordinated process responding to multiple ligands within the injury niche. Furthermore, the use of spatial transcriptomics in pulmonary fibrosis patients reinforced the potential role for inflammatory fibroblasts in injury repair and disease. In these fibrotic lungs inflammatory fibroblasts were found in injury sites (**Figure 7**), away from the airways and closer to regions of alveolar remodeling. The enrichment of these cells in these sites indicates that even in late-stage disease these fibroblasts are participating in the injury response, however the implications that these cells are not close to the pathological KRT5-/KRT17+ cell is an important observation that requires further study.

We must note limitations to this study that should be considered when framing the interpretation of our findings. First, the functional biology of the inflammatory fibroblast was performed in murine injury. Though our findings are consistent with the established literature in the mouse, the human lung contains anatomical differences and additional cell types not seen in the mouse lung. Additionally, a strength of mouse models is that they allow for temporal modeling of lung injury prior complete loss of regenerative capacity, however human lung samples are typically collected from patients immediately prior to transplantation and have likely lost the critical niches necessary for appropriate repair. Also, we rely on reductionist organoid models to describe the interaction of epithelial and fibroblast populations, however this does not consider the role of inflammation and immune cells in the injury repair process. Though we did not see an association between increased inflammatory fibroblasts and sustained inflammation in our injury model, this does not rule out activation of immune cells as a critical component of inflammatory fibroblast function. Moreover, the lack of a BMP inhibitory signature in human transcriptomic data and the sustained inflammatory cytokine signature in this population suggests immune crosstalk is likely. Future studies focusing on this aspect of the inflammatory fibroblast are planned.

In summary our findings are an important step forward in a coordinated effort by our field to understand the implication of fibroblast heterogeneity across lung injury. We show that there is remarkable consistency across injury modalities and fibroblast response. Our identification of BMP4 from inflammatory fibroblasts complements observations described in the literature for more than 20 years and suggests that the process of wound healing is linked closely to developmental pathways. Finally, the observed presence of these fibroblasts across multiple human diseases prompts important questions to consider as we move forward with therapeutic approaches to resolve acute and chronic injury in patient populations.

## METHODS

### Sftpc^I73T^ Mouse Model of Pulmonary Fibrosis

Tamoxifen inducible *Sftpc*^I73T/I73T^ Rosa26ERT2FlpO^+/+^ (a.k.a. I^ER^-*Sftpc^I73T^*) mice expressing an NH_2_-terminal HA-tagged murine *Sftpc^I73T^* mutant allele into the endogenous mouse *Sftpc* locus were previously generated as reported^30^ and are summarily detailed. Tamoxifen treatment (360 mg/kg in males and 520 mg/kg in females) of adult I^ER^-*Sftpc^I73T^* mice was initiated at 12-14 weeks of age by oral gavage (OG) split evenly on day 0 and day 3. All mouse strains and genotypes generated for these studies were congenic with C57/B6/J. Both male and female animals (aged 8-14 weeks) were utilized in tamoxifen induction protocols. All mice were housed under pathogen free conditions in an AALAC approved barrier facility at the Perelman School of Medicine, University of Pennsylvania. All experiments were approved by the Institutional Animal Care and Use Committee at the University of Pennsylvania.

### Reagents and Materials

Cytological stains used were Diff-Quik® (Thermo Fisher Scientific, Inc., Pittsburgh, PA) and Giemsa (#GS500; MilliporeSigma (St. Louis, MO)). Tamoxifen (non-pharmaceutical grade) was purchased from MilliporeSigma. Except where noted, all other reagents were electrophoretic or immunological grade and purchased from either MilliporeSigma or Thermo-Fisher. Antibodies-Antibodies used for flow-cytometry and FACS were obtained from commercial sources (**Supplemental Table 1**).

### Multichannel Flow Cytometry for Identification of Lung Cell Populations

Previously validated sorting strategies ^20,22,30,42,69–71^ were applied in the following manner. Blood free perfused lungs were digested in Phosphate Buffered Saline (Mg and Ca free) with Collagenase Type I (Gibco Cat# 17100017), DNase (Millipore Sigma Cat# D5025), and Dispase (BD Biosciences). Resulting product was passed through 70 μm nylon mesh to obtain single-cell suspensions and then processed with ACK Lysis Buffer (Thermo Fisher). Cells were incubated with antibody mixtures (see **Supplemental Table 1**). Stained cells were analyzed with an LSR Fortessa (BD Biociences). AT2 cells were identified as EpCAM+, CD45-, CD31-, MHCII^hi^, CD104-. Fibroblasts were identified as EpCAM-, CD45-, CD31-, CD140a+. With some adaptions of previously reported strategies ^20,22,42^. Specifically, alveolar fibroblasts were defined as PDGFRA+, MCAM-, Sca1-, adventitial fibroblasts were defined as PDGFRa+, MCAM-, Sca1+, fibrotic fibroblasts were defined as PDGFRA+, MCAM-, Sca1-, CD9+, inflammatory fibroblasts were derived from injured lungs and defined as PDGFRA+, MCAM-, Sca1-. This data was analyzed with FlowJo software (FlowJo, LLC, Ashland, Oregon).

### Lung Histology, H&E severity, and Immunofluorescence Staining

Whole lungs were fixed by tracheal instillation of 10% neutral buffered formalin (MilliporeSigma) at a constant pressure of 25 cm H_2_O. 6 ◻m sections were stained with Hematoxylin & Eosin (H&E) or Masson’s Trichrome stains by the Pathology Core Laboratory of Children’s Hospital of Philadelphia. Slides were scanned using an Aperio ScanScope Model: CS2 (Leica) at 10X magnification and representative areas captured and exported as TIF files and processed in Adobe Illustrator. Whole lung lobe Trichrome images were color coded blue (normal), green (moderate), and severe (red), and scored for fibrosis severity as the percent of total lung lobe area using a published algorithm^62^. Staining of lung sections for fibrillar collagen was performed using the Picrosirius Red (PSR) Stain Kit following the manufacturer’s instructions (Polysciences, Inc., Warrington PA). Digital morphometric measurements were performed on multiple lobes and multiple levels with ten random peripheral lung images devoid of large airways per slide analyzed at a final magnification of 100× using Image J as published (34, 45, 46). The mean area of each lung field in each section staining for PSR was calculated and expressed as a percentage of total section area as adapted from Henderson, et al (47). Immunofluorescence chemistry (IFC) staining was performed on 6 μm paraffin embedded lung sections using a combination of commercially available primary antibodies and fluorescent AlexaFluor anti-IgG secondary antibodies (Supplementary Table 1). Images were obtained using an Eclipse Ti2 Series inverted microscope (Nikon) and analyzed using Nikon NES-Elements software and Fiji 2.15.1.

### Bronchoalveolar Lavage Fluid (BALF) Collection, Processing, and Cytokine Measurement

BALF collected from mice using sequential lavages of lungs with five X 1 ml aliquots of sterile saline was processed for analysis as previously described^30^ and detailed in the supplemental methods. Total acid soluble collagen content in cell free BALF was determined using the Sircol assay kit (Biocolor, Ltd; Carrickfergus, UK) according to the manufacturer’s instructions.

### RNA Isolation and Quantitative Real Time Polymerase Chain Reaction

RNA was extracted from homogenized lung suspension using RNeasy Mini Kit (Qiagen, Valencia, CA) following the manufacturer’s protocol. The concentration and quality of extracted RNA from the lung tissues were measured using NanoDrop® One (Thermo Scientific, Wilmington, DE) and reverse-transcribed into cDNA using either Taqman Reverse Transcription Reagents (Applied Biosystems, Foster City, CA) or Verso cDNA Synthesis Kit (ThermoFisher). Quantitative real time PCR was performed using Sybr Green qPCR master mix (ThermoFisher) on a QuantStudio 7 Flex Real-Time PCR System with results normalized to 18S and Actb RNA. Primer sequences for all mouse genes are listed in **Supplemental Table 1**.

### Measurement of Pulmonary Function

At takedown, mice underwent Flexivent (SCIREQ, Inc. Toronto Canada) analysis for assessment of lung physiology as previously described^72,73^. Briefly, invasive measurement of static lung compliance was performed with mice anesthetized with intraperitoneal pentobarbital. The mouse tracheas were cannulated with a 20-gauge metal stub adapter and then placed on a small-animal ventilator at 150 breaths per min and a tidal volume of 10 ml/kg of body weight. Static lung compliance was determined with the manufacturer’s software using a 2 second breath pause maneuver.

### Population RNA Sequencing

Isolated RNA from sorted fibroblast populations was sent for library preparation by GeneWiz, LLC. Fastq files were processed as previously described^71^ and detailed in the supplemental methods. Heatmaps were generated using Morpheus (https://software.broadinstitute.org/morpheus). Protein-protein interactions were obtained using STRING (Search Tool for Retrieval of Interacting Genes/Proteins) database. Gene ontogeny analysis was performed using the Database for Annotation, Visualization, and Integrated Discovery (https://david.ncifcrf.gov/home.jsp) based on differentially enriched genes (FC >1.5 P-value <0.05). Key pathway analyses were performed on gene lists identified from the GSEA molecular signature database^74–76^.

### Single cell RNA sequencing and Analysis of Lung Cell Populations

To capture representative proportions of epithelial and mesenchymal populations, we profiled 105,340 cells from the *Sftpc*^I73T^ model, this includes previously published data sets (GSE234604 and GSE296513) as well as newly generated data for this study. The new single cell suspensions were prepared by physical and enzymatic dissociation followed by Dynabeads™ M-280 Streptavidin (Invitrogen™ 11205D) depletion of CD45+ and CD31+ cells using CD45 and CD31 biotinylated antibodies (**Supplemental Table 1**). Cell suspensions were then loaded onto individual GemCode instrument (10X Genomics). Single-cell barcoded droplets were produced using 10X Single Cell 3’ v3 chemistry. Libraries generated were sequenced using the HiSeq Rapid SBS kit, and the resulting libraries were sequenced across the two lanes of an Illumina HiSeq2500 instrument in a High Output mode. Single cell RNA-Seq reads were aligned to mouse genome (mm10/GRCm38) using STARSolo (version 2.7.5b). After initial quality control and processing, we analyzed the scRNA-seq data using the Scanpy pipeline (50). Genes expressed in fewer than 3 cells were removed, and cells with fewer than 200 genes and a mitochondrial fraction of less than 20% were excluded. Counts were log-normalized using scanpy.pp.normalize_per_cell (counts_per_cell_after=1×104), followed by scanpy.pp.log1p. To integrate data from multiple samples, we used Scvi-tools (51). We applied scvi.model.SCVI.setup_anndata() to establish the model parameters for integration, including: layer, categorical_covariate_keys, and continuous_covariate_keys. We then performed a principal component analysis (PCA) and generated a K-nearest neighbor (KNN) graph using scanpy.pp.neighbors with n_neighbors=15. The resulting KNN graph was used to perform Uniform Manifold Approximation and Projection (UMAP) dimension reduction to visualize the cells in two dimensions using scanpy.tl.umap(). Clustering was performed using the Leiden algorithm with scanpy.tl.leiden (52). We identified cell populations using known canonical marker genes or by assessing cluster-defining genes based on differential expressions.

Integration of published murine data sets was performed in a similar approach using Scanpy and Scvi-tools. First all data sets were obtained from GSEs listed in data availability and combined into a single adata object. Some files were obtained as H5ads with previous annotations, when possible, we generated our objects from the individual matrix, barcodes, and genes files. The combined object was processed and filtered as above followed by integration via Scvi-tools again as described above. The mesenchyme was identified as negative for Epcam, Pecam1, and Ptprc with additional filtering out of small clusters containing more than 25 ribosomal marker genes in the top 50 markers genes defined by sc.tl.rank_genes_groups().

Analysis of the integrated Human Lung Cell Atlas ^77^ was performed using the annotated file labeled as HLCA Full (https://data.humancellatlas.org/hca-bio-networks/lung/atlases/lung-v1-0). This annotated H5ad has a “Stroma” annotation (ann_level_1) that was used to define the mesenchymal cell subset (181,028 cells) analyzed further in this manuscript. This dataset was made up of 23 different studies sequenced using 8 different assays. We further limited the analysis to the two assays that contained the most cells (10x 3’ v2 and 10×5’ v1) for a final stromal object of 79,549 cell. As before we performed additional filtering of small clusters containing more than 25 ribosomal maker genes in the top 50, followed by cluster identification using the markers defined in in **Figure 6**.

### Primary Mouse Organoid Culture

Organoid culture assays were performed as previously described^70^. AT2 cells and fibroblasts were flow sorted as described above. In each technical replicate, 5000 AT2 cells were combined with 50,000 *Pdgfra*^+^ lung fibroblasts in 50% Matrigel (Corning) and 50% SAGM (Lonza) in a Falcon Cell Culture Insert. Cell/matrigel suspension solidified and SAGM medium was then added into the bottom of the well. SAGM was prepared according the BulletKit (Lonza) manufacturer instructions with some modification (Hydrocortisone, BSA, Triiodothyronine, and Epinephrine were not included). Medium was changed every other day. 10 μM rock inhibitor (Y-27632 dihydrochloride, Millpore Sigma, catalog # Y0503) was added to the medium for the first two days. AT2 alone organoid culture was also performed as previously described^24,78^. Freshly flow sorted AT2 cells were resuspended at GFR-Matrigel at 100 cells/μL and 50 μL Matrigel droplets were plated in a 12-well plate (Falcon). Matrigel droplets were allowed to solidify for 30 minutes at 37°C. For media, previously published AT2 Maintenance Media (AMM)^78^ and minimally modified. AMM was composed of 10 μM SB431542 (Abcam), 3 μM CHIR99021 (Tocris), 1x Insulin-Transferrin-Selenium (ITS) (Thermo), 15mM HEPES (Thermo), 1x Glutamax (Thermo), 50ng/mL hEGF (Gibco), 5μg/mL Heparin (Sigma), 1.25mM N-Acetyl Cysteine (Sigma), and 10ng/ml FGF7 (Tocris) in Advanced DMEM/F12 (Gibco). For the first two days of culture AMM was additionally supplemented with 10 μM rock inhibitor (Y-27632 dihydrochloride, Sigma). Subsequently, AMM was refreshed every other day for the duration of the experiment. Organoids were imaged on an EVOS FL Microscope and were quantified via ImageJ using the “analyze particles” macro.

### Spatial Transcriptomic Analysis of Pulmonary Fibrosis and Control Lung Tissue

We reanalyzed data that was previously obtained and published (GSE250346) from patients with pulmonary fibrosis and unaffected lung tissue. Tissues were previously assigned disease status (healthy donor ‘HD’, interstitial lung disease ‘ILD’) and additionally labeled as either less affected (<75% pathology) or more affected (>75% pathology) as determined by percent pathology scores^14^. A total of 45 samples were analyzed with 27 histological features including 5 mesenchymal/interstitial (interlobular septum, airway smooth muscle, fibroblastic focus, fibrosis and severe fibrosis). Description of methodology for identification of cell-based niches is fully described in the previous publication and those same cell-based niches were applied here. To match the inflammatory fibroblasts population in the spatial data with that of the HLCA we expanded the existing annotated inflammatory fibroblasts population to include cells within the mesenchymal lineage with detectable expression (raw count ≥ 1) of at least two of the following markers: CCL2, PTGDS, and HAS2. All other cell type annotations were retained for this work. Enrichment of inflammatory fibroblasts within cell-based niches and pathology annotated regions was assessed using fisher exact tests. P-values were adjusted using Benhamini-Hochberg false discover rate approach.

To assess proximity between inflammatory fibroblasts and other cell types, a cell proximity analysis was performed on spatially resolved single-cell data. For each sample, cell coordinates were used to construct a spatial point pattern, and all cell pairs within a distance threshold of 30 µm were identified using the *closepairs* function from the *spatstat* R package. Pairwise distances and angles were computed, with angular space divided into 30° bins to capture the nearest neighbor in each angular direction. Only first-degree neighbors within angular directions were considered proximal to the anchored cell. The probability of cell types being proximal to one another was assessed using logistic regression by assigning cells to binarized proximal and nonproximal categories and using cell type as a covariate. Odds ratios and 95% confidence intervals were calculated relative to the mean predicted log-odds across all cell types. The analysis was performed both across all samples and within niches.

### Statistics

All data are presented with dot-plots and group mean ± SEM unless otherwise indicated. Statistical analyses were performed with GraphPad Prism (San Diego, CA). Two tailed Student’s t-test were used for 2 groups as indicated; Multiple comparisons were performed by analysis of variance (ANOVA) with post hoc testing as indicated; survival analyses was performed using Kaplan Meier with Mantel Cox correction. In all cases statistical significance was considered at p values ≤ 0.05.

## Data and code availability

The sequencing data generated in this study will be made available in GEO upon publication. The HLCA was downloaded from cellxgene (https://cellxgene.cziscience.com/collections/6f6d381a-7701-4781-935c-db10d30de293). Analysis of previously published data was performed on GSE253453, GSE221402, GSE132771, GSE234604, GSE296513, GSE215094, GSE248798, GSE244237, GSE249931, GSE229523, GSE276546, GSM6428697.

## Study Approval

Mice housed in pathogen free facilities were subjected to experimental protocols approved by the IACUC of the Perelman School of Medicine at the University of Pennsylvania.

## AUTHOR CONTRIBUTIONS

LRR and MFB developed the concept. LRR, WRB, and MFB designed the experiments. YT, CHC, AR, and RC performed *in vivo* animal experiments. LRR, WRB, AM, NH, SB, DLJ, CHC, AR, RC, ETH, SM, CHC conducted experiments and analyzed data. LRR and WRB performed bioinformatic analysis. LRR and AM modified, validated, and optimized flow cytometry strategies. LRR, WRB, NEB, ETH, JK, JAK, NEB, and MFB interpreted data and generated figures. LRR and WRB drafted the original manuscript. LRR, WRB, AM, SB, JAK, NEB, and MFB edited the manuscript. All authors reviewed and approved the final version prior to submission.

## ACKNOWLEDGMENTS

The authors wish to thank all members of the Beers and Katzen labs for insightful discussions. Michael F. Beers is the Robert L. Mayock and David A. Cooper Professor of Medicine. The authors thank the Penn Genomic and Sequencing Core (RRID:SCR_022383). The data for this (manuscript) were generated in the Penn Cytomics and Cell Sorting Shared Resource Laboratory at the University of Pennsylvania (RRID:SCR_022376). Penn Cytomics is partially supported by the Abramson Cancer Center NCI Grant(P30016520). We thank the PENN-CHOP Lung Biology Institute Informatics team for guidance. Multiple figures were created using Biorender.com.

