## Supplemental Data and Methods for "Inflammatory Fibroblasts Promote Repair After Injury Through Epithelial Proliferation"

**¶ M.F.B. is the corresponding author**

**Running Title:** *Inflammatory Fibroblasts in Lung Injury and Repair*

**Lead Contact Author:**

Michael F. Beers, M.D.
Pulmonary and Critical Care Division
Perelman School of Medicine at The University of Pennsylvania
Edward J Stemmler Hall Suite 216
3450 Hamilton Walk
Philadelphia, Pennsylvania 19104-6118


**Conflict of Interest:** The authors declare that no conflicts of interest exist.

### SUPPLEMENTAL METHODS

##### *Bronchoalveolar Lavage Fluid (BALF) Collection, Processing, and Cytokine Measurement*

BALF collected from mice using sequential lavages of lungs with five X 1 ml aliquots of sterile saline was processed for analysis as previously described[1]. First ml of saline wash is labeled as enriched cell free BALF used to measure total protein and cytokines. Cell pellets obtained by centrifuging BALF samples at 400 g for 6 minutes were re-suspended in 1 ml of PBS, and total cell counts determined using a NucleoCounter (New Brunswick Scientific, Edison, NJ). Differential cell counts were determined manually from BALF cytospins stained with modified Giemsa (Sigma Aldrich, #GS500) to identify macrophages, lymphocytes, eosinophils and neutrophils. Total protein content of cell free BALF was determined using the DC Protein Assay Kit (Cat # 5000111; BioRAD, Inc, Hercules CA) with bovine serum albumin as a standard according to the manufacturer's instructions.

##### *RNA Isolation and Reverse Transcriptase Quantitative Polymerase Chain Reaction (RT-qPCR)*

RNA was extracted by first lysing cells in Qiazol (Qiagen) and subsequently using RNAeasy mini kit (Qiagen) according to the manufacturer's protocol. The concentration and quality of extracted RNA from the lung tissues were measured using the NanoDrop® (Thermo Scientific, Wilmington, DE) and reverse-transcribed into cDNA using Verso cDNA Synthesis Kit (ThermoFisher). RT-qPCR was performed on a QuantStudio 7 Flex Real-Time PCR System with results normalized to *18S* and *Actb* gene expression. Primer sequences in **Supplemental Table 1**.

##### *Population RNA Sequencing Data Processing*

Fastq files were evaluated for quality control with the FastQC program and then aligned against the mouse reference genome (mm10) using the STAR aligner[2]. Duplicate reads were flagged with the MarkDuplicates program from Picard tools and excluded from analysis. Per gene read counts for Ensembl (v67) gene annotations were computed using the R package Rsubread. Gene counts, represented as counts per million (CPM), were nominalized using TMM method in the edgeR R package, and genes with 25% of samples with a CPM < 1 were considered low expressed and removed. The data were transformed with the VOOM function from the limma R package to generate a linear model and perform differential gene expression analysis[3]. We employed the empirical Bayes procedure as implemented in limma to adjust the linear fit and to calculate P values given the small sample size of the experiment. We adjusted P values for multiple comparisons using the Benjamini-Hochberg procedure. Heatmaps were generated using the Broad Institute's online tool Morpheus ( https://software.broadinstitute.org/morpheus). Protein-protein interactions were obtained using STRING (Search Tool for Retrieval of Interacting Genes/Proteins) database. Gene ontogeny analysis was performed using the Database for Annotation, Visualization, and Integrated Discovery (https://david.ncifcrf.gov/home.jsp) based on differentially enriched genes (FC >1.5 P-value <0.05). Key pathway analyses were performed on gene lists identified from the GSEA molecular signature database[4–6].

| Supplemental Table 1: Antibodies and Primer Sequences | |
|---|---|
| Antibodies | |
| mKi67 (28074-1-AP) | Proteintech |
| HA Tag Monoclonal (81290-1-RR) | Proteintech |
| Ly6C (HK1.4) | Biolegend |
| Mcam (ME-9F1) | Biolegend |
| CD9 (KMC8) | Biolegend |
| CD11b (M1/70) | BD Biosciences |
| CD11c (HL3) | Biolegend |
| Ly6G (1A8) | Biolegend |
| CD64 (X54-5/7.1) | Biolegend |
| CD43 (S11) | Biolegend |
| CD3e (KMC8) | BD Biosciences |
| Foxp3 (FJK-16s) | Thermofisher |
| CD45 (30F-11) | Biolegend |

| | |
|---|---|
| CD25 (PC61) | BD Biosciences |
| Epcam (G8.8) | Biolegend |
| Epcam (G8.8) | BD Biosciences |
| CD31 (MEC13.3) | Biolegend |
| CD104 (346-11A) | Biolegend |
| CD51 (RMV-7 | Biolegend |
| I-A/I-E,MHCII (M5/114.15.2) | Biolegend |
| CD140a/PDGFRa (APA5) | Biolegend |

| Primer Sequences | | |
|---|---|---|
| Target | Forward | Reverse |
| Sfrp1 | CCTCTAAGCCCCAAGGTACA | GACTGGAAGGTGGGACACTC |
| Hp | TCCACGATGAGAGCCCTGG | CATCCATAGAGCCACCGATGA |
| Ces1d | CCCATTGCTGGTCTGGTTGC | TGCCTTCAGCGAGTGGATAG |
| Inmt | AGCCTGCAGAACCTCTACCA | TGCCACCTGCTTCTGTCTCC |
| Slc7a10 | GCACCATCATCATCGGGAAC | AGCACTCCAGAGCAGTAGGA |
| Ebf1 | TCTATGTGCGCCTCATCGAC | AGGGAGTAGCTGCATGTTCC |
| 18S | CGGCTACCACATCCAAGGAA | GCTGGAATTACCGCGGCT |
| Cthrc1 | ATCCCAGGTCGGGATGGATT | CCAATCCCTTCACAGAGTCCT |
| Col14a1 | TGAAGCACCCACAGCCATAG | TCCAGGCACCATAACCGTTC |
| Col13a1 | CTAAAGGGGAGATGGGCCTG | ATTATCGGGTTGGAGCGCAG |
| Pi16 | AACTATGAGCCTCCGGGGAA | GTCACCCTTGGAGGCGAATC |
| Plin2 | GCTCTCCTGTTAGGCGTCTC | AACAATCTCGGACGTTGGCT |
| Npnt | AGCATGTATCCGTTGAGACAGTA | GAAGCCTCGGCCCTGTAAC |
| Pdgfra | CTGTTGGAGCTTGAGGGAGAG | ATAGCTCCTGAGACCTTCTCCT |
| Lcn2 | CTGTACCTGAGGATACCTGTGC | TGAGTGTCATGTGTCTGGGC |
| Saa3 | CTCGCAGCACGAGCAGGAT | GCATCATAGTTCCCCCGAGC |
| Hp | TCCACGATGAGAGCCCTGG | CATCCATAGAGCCACCGATGA |
| Spp1 | AGCAAGAAACTCTTCCAAGCAA | GTGAGATTCGTCAGATTCATCCG |

**Figure S1**

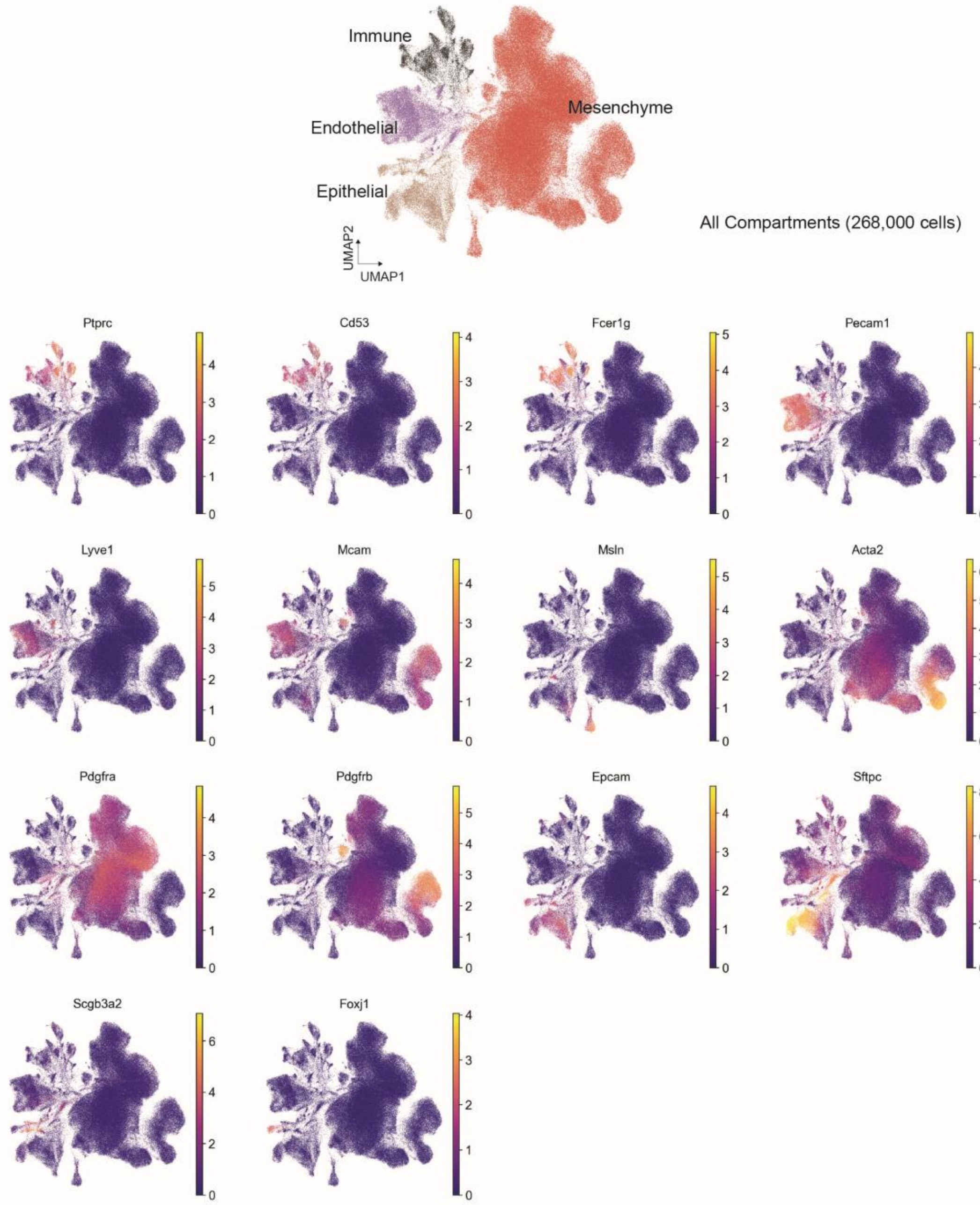
Immune
Endothelial
Epithelial
Mesenchyme
All Compartments (268,000 cells)
UMAP2
UMAP1
Ptprc
Cd53
Fcer1g
Pecam1
Lyve1
Mcam
Msln
Acta2
Pdgfra
Pdgfrb
Epcam
Sftpc
Scgb3a2
Foxj1

**Figure S1. Integration of multiple single cell data sets identifies four major cellular compartments.** UMAP projection of the integrated dataset described in Figure 1 color-coded by major cellular compartments. UMAP projection of the integrated dataset color-coded by major compartment identifying genes.

**Figure S2**

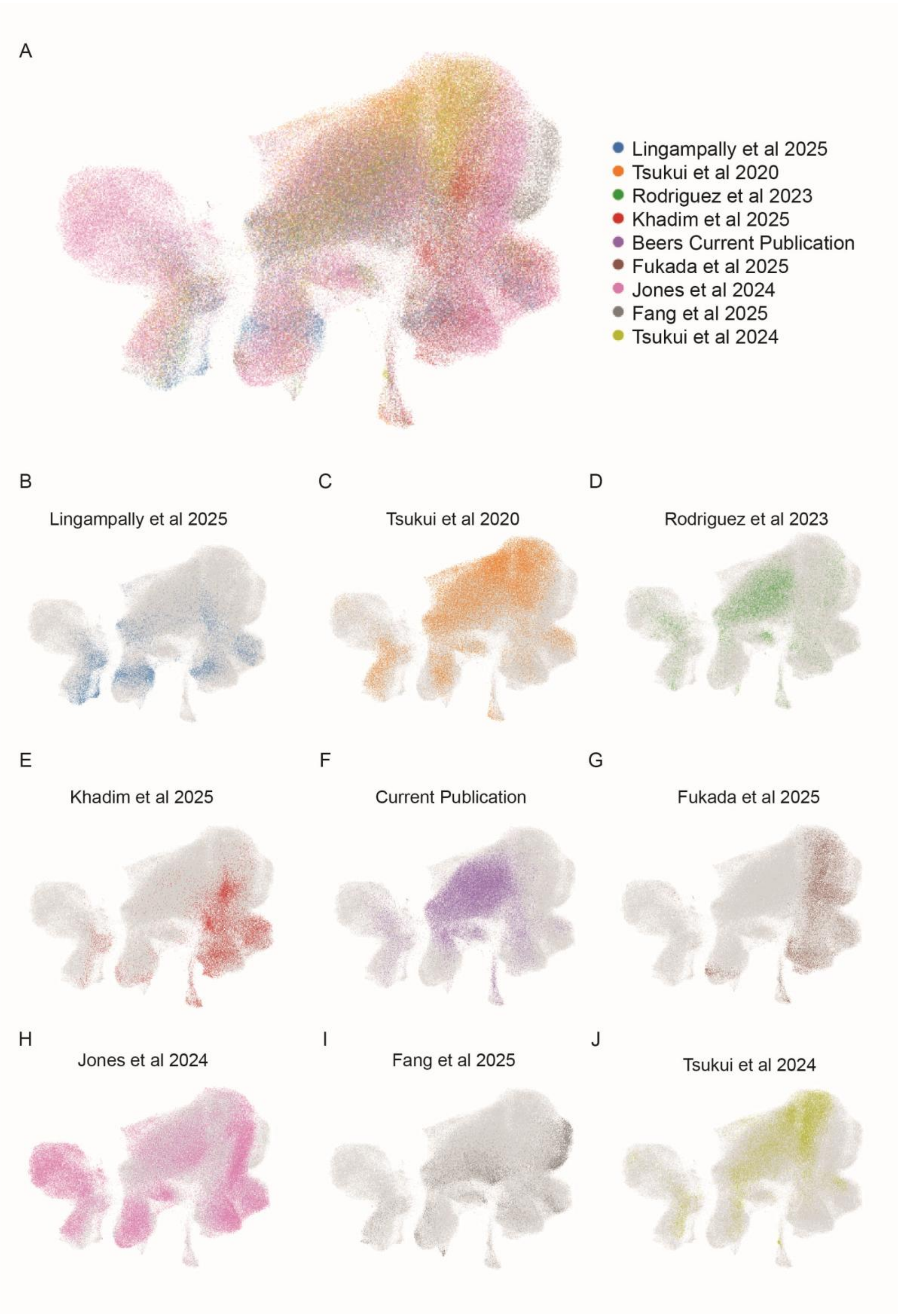
A
Lingampally et al 2025
Tsukui et al 2020
Rodriguez et al 2023
Khadim et al 2025
Beers Current Publication
Fukada et al 2025
Jones et al 2024
Fang et al 2025
Tsukui et al 2024
B
Lingampally et al 2025
C
Tsukui et al 2020
D
Rodriguez et al 2023
E
Khadim et al 2025
F
Current Publication
G
Fukada et al 2025
H
Jones et al 2024
I
Fang et al 2025
J
Tsukui et al 2024

**Figure S2. Integration of multiple single cell data sets confirms population identities are conserved across all studies**. A)**.** UMAP projection of the integrated dataset described in Figure 1 color-coded by study IDs B) UMAP projection of the integrated dataset color-coded by major compartment study ID, demonstrating distribution of each data set.

**Figure S3**

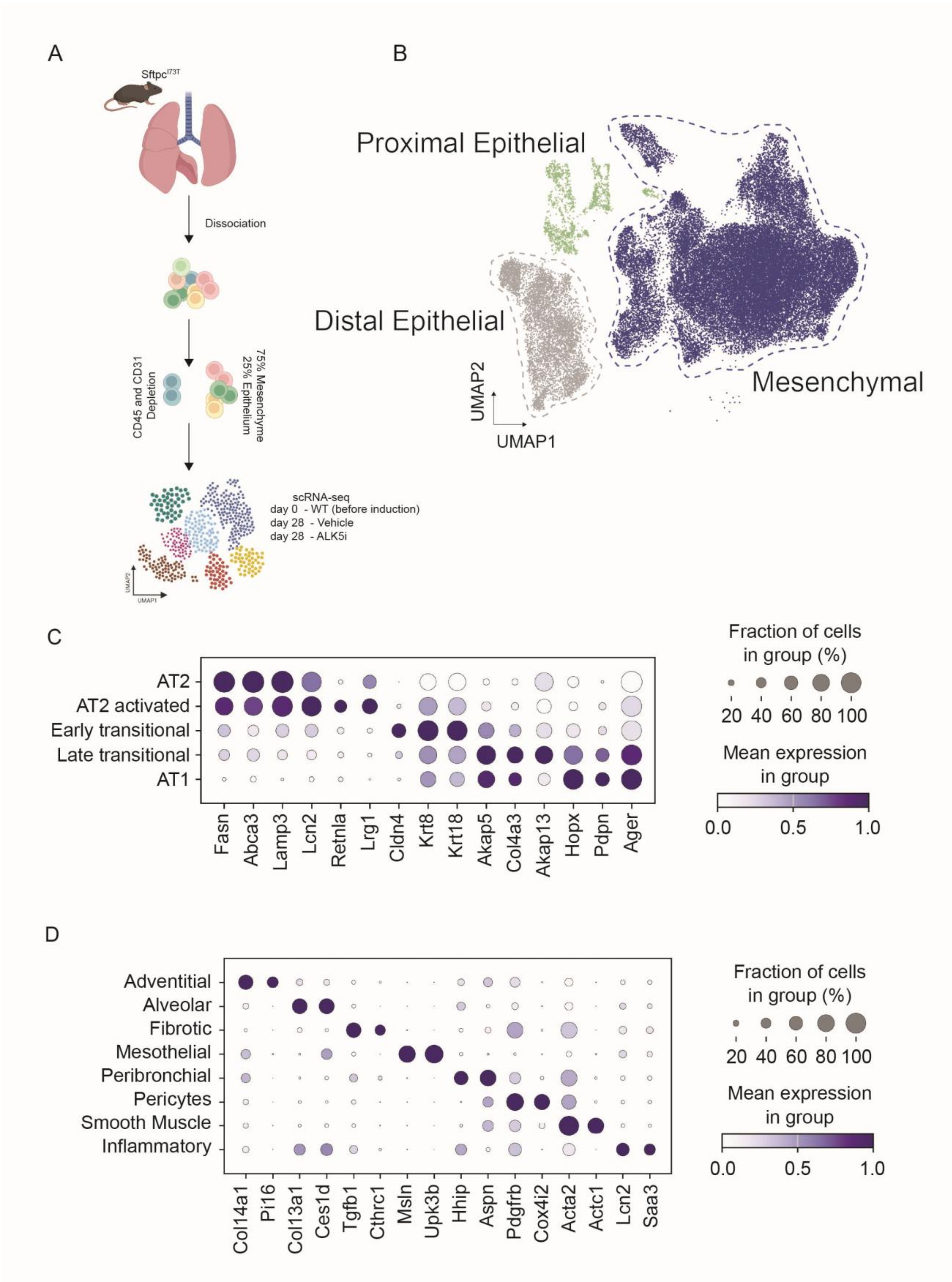
A
Sftpc I73T
Dissociation
CD45 and CD31 Depletion
75% Mesenchyme
25% Epithelium
scRNA-seq
day 0 - WT (before induction)
day 28 - Vehicle
day 28 - ALK5i
B
Proximal Epithelial
Distal Epithelial
Mesenchymal
UMAP2
UMAP1
C
AT2
AT2 activated
Early transitional
Late transitional
AT1
Fasn
Abca3
Lamp3
Lcn2
Retnla
Lrg1
Cldn4
Krt8
Krt18
Akap5
Col4a3
Akap13
Hopx
Pdpn
Ager
Fraction of cells in group (%)
20 40 60 80 100
Mean expression in group
0.0 0.5 1.0
D
Adventitial
Alveolar
Fibrotic
Mesothelial
Peribronchial
Pericytes
Smooth Muscle
Inflammatory
Col14a1
Pi16
Col13a1
Ces1d
Tgfb1
Cthrc1
Msln
Upk3b
Hhip
Aspn
Pdgfrb
Cox4i2
Acta2
Actc1
Lcn2
Saa3
Fraction of cells in group (%)
20 40 60 80 100
Mean expression in group
0.0 0.5 1.0

**Figure S3. Cluster defining markers in epithelial and mesenchymal enrichment of Sftpc$^{I73T}$ single cell data.** A) Schematic representation of the scRNA-seq strategy used to generate the *Sftpc*$^{I73T}$ datasets, which involved depletion of CD45+ and CD31+ cells to achieve an approximate ratio of 25% epithelial and 75% mesenchymal cells. B) UMAP projection of the integrated data set highlighting the three compartments analyzed C-D) Frequency and expression analysis using dot plots to highlight marker genes used to define each epithelial (C) and mesenchymal cluster (D).

**Figure S4**

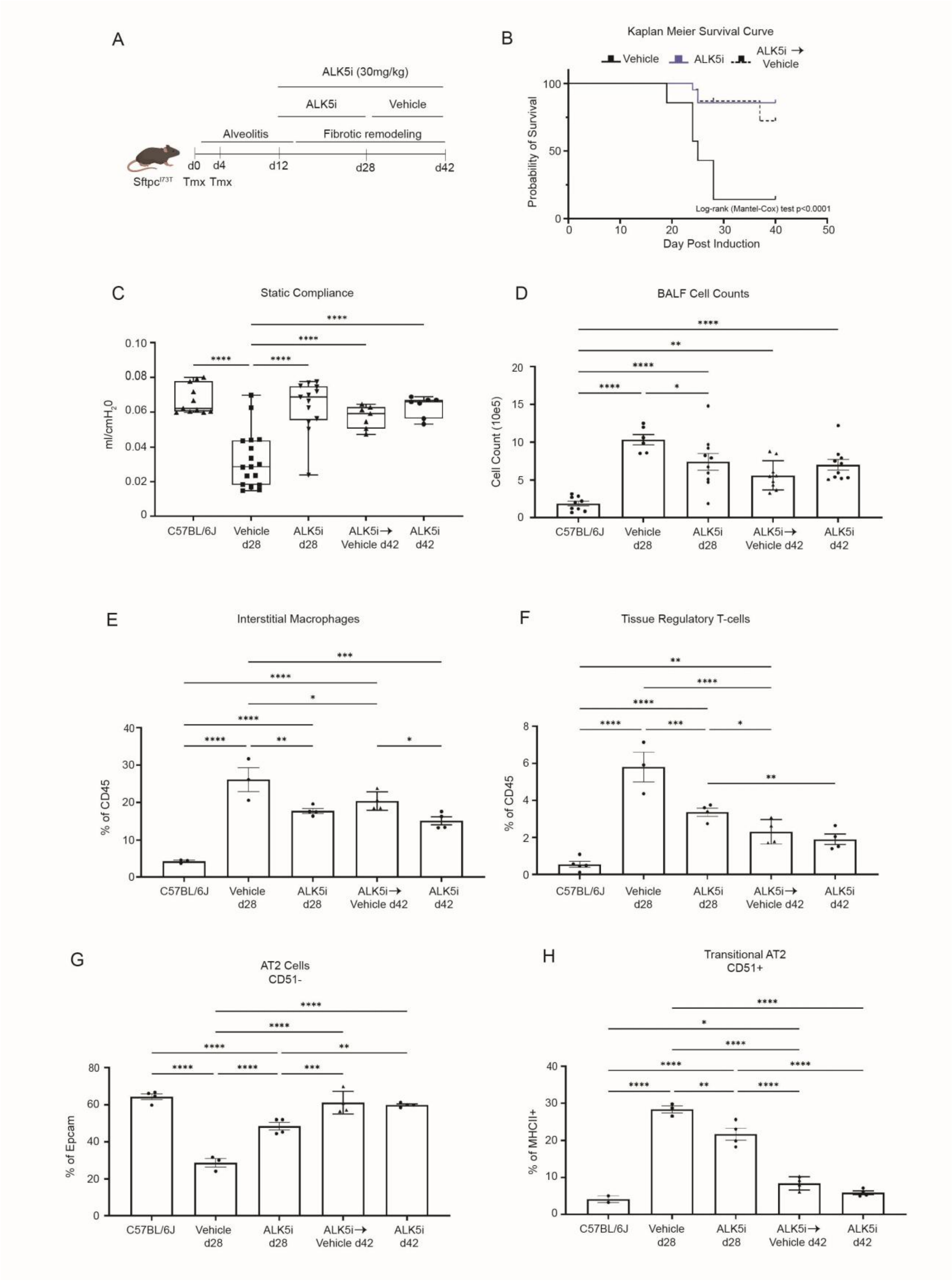
A
ALK5i (30mg/kg)
ALK5i
Vehicle
Alveolitis
Fibrotic remodeling
d0 d4 d12 d28 d42
Sftpc^I73T Tmx Tmx
B
Kaplan Meier Survival Curve
Vehicle ALK5i ALK5i → Vehicle
Probability of Survival
Day Post Induction
Log-rank (Mantel-Cox) test p<0.0001
C
Static Compliance
ml/cmH2O
C57BL/6J
Vehicle d28
ALK5i d28
ALK5i→ Vehicle d42
ALK5i d42
D
BALF Cell Counts
Cell Count (10e5)
E
Interstitial Macrophages
% of CD45
F
Tissue Regulatory T-cells
% of CD45
G
AT2 Cells CD51-
% of Epcam
H
Transitional AT2 CD51+
% of MHCII+

**Figure S4. Extended TGFB inhibition does not promote inflammation in *Sftpc*$^{I73T}$ mice** A) Schematic of intervention protocol outlining tamoxifen delivery to induce expression of SPC-C$^{I73T}$ at days 0 and 4 followed by daily intervention with oral gavage (o.g.) ALK5 inhibitor SB525334 (60 mg/kg BID, n = 20 mice per group) until day 28 where the ALK5 inhibitor group was further divided into two subsets. Ones subgroup continued the intervention for 14 days while the other received vehicle control. B) 28-day mortality was significantly reduced in ALK5i group but extended 42-day mortality did not change between subgroups.C) Static lung compliance, measured by Scirec Flexivent, increased significantly in all ALK5i groups compared to vehicle control (mean ± SEM; n=7-17 all groups). D) Total cell counts bronchoalveolar lavage (BAL) fluid collected from mice at 28d or 42d post tamoxifen were significantly reduced in all ALK5i-treated mice. E-F) Flow cytometry quantification of whole lung single cell suspension demonstrates decreased accumulation of fibrosis associated macrophages and T regulatory cells after ALK5 intervention, these populations are unchanged after ALK5 is removed (mean ± SEM; n=3-5 mice per group). G-H) Flow cytometry quantification of whole lung single cell suspension demonstrates the rescue of AT2 cell numbers by ALK5 inhibition and a decrease in AT2-AT1 transitional cell numbers at 28 days post induction with further improvement towards control conditions at 42 days post induction regardless of ALK5i subgroups (mean ± SEM; n=3-5 mice per group).

**Figure S5**

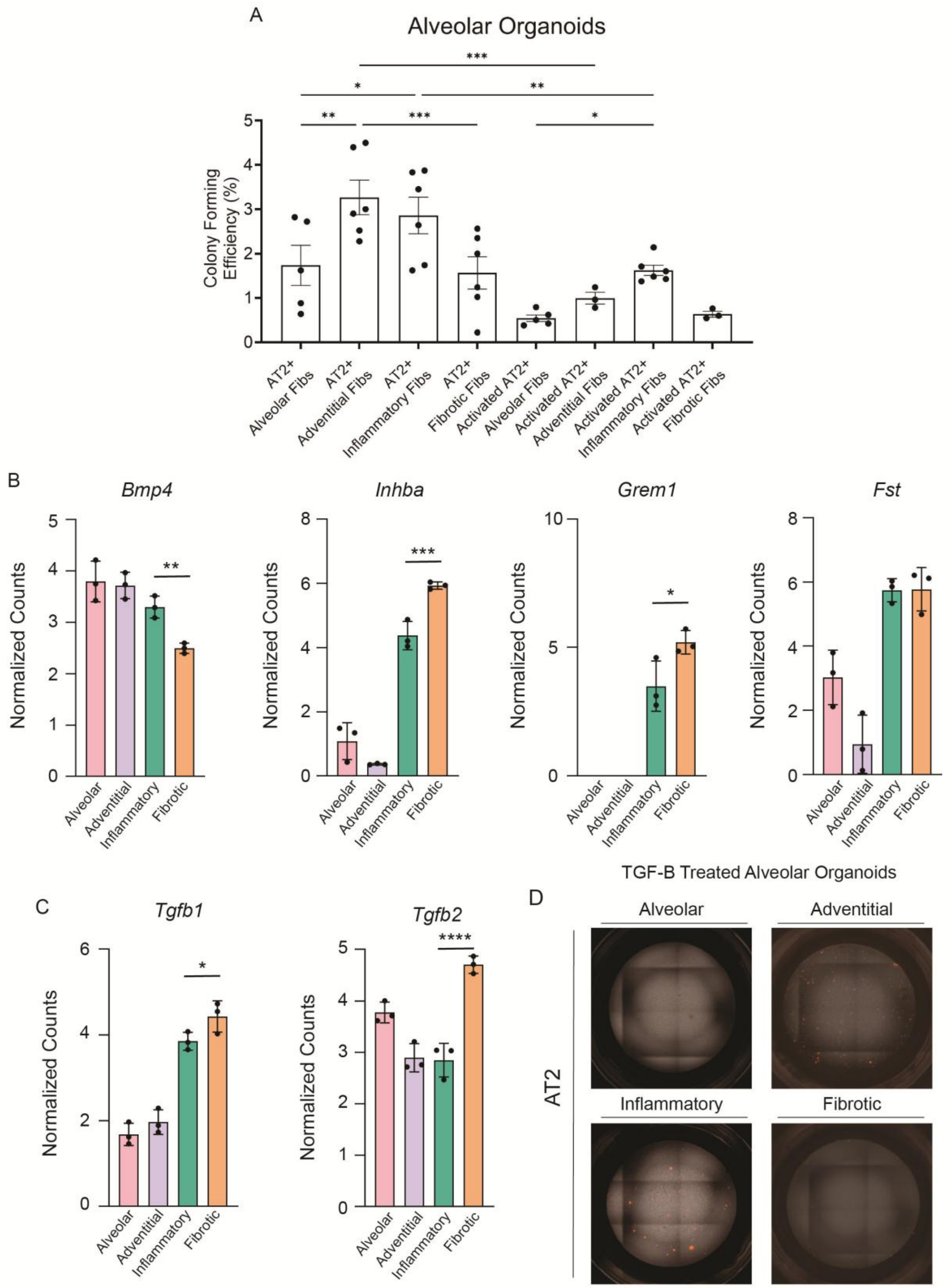
A
Alveolar Organoids
Colony Forming Efficiency (%)
AT2+ Alveolar Fibs
AT2+ Adventitial Fibs
AT2+ Inflammatory Fibs
AT2+ Fibrotic Fibs
Activated AT2+ Alveolar Fibs
Activated AT2+ Adventitial Fibs
Activated AT2+ Inflammatory Fibs
Activated AT2+ Fibrotic Fibs
B
Bmp4
Inhba
Grem1
Fst
Normalized Counts
Alveolar
Adventitial
Inflammatory
Fibrotic
C
Tgfb1
Tgfb2
D
TGF-B Treated Alveolar Organoids
Alveolar
Adventitial
Inflammatory
Fibrotic
AT2

**Figure S5. Decreased BMP inhibition in fibroblasts after lung injury.** A) Colony forming efficiency (CFE; %) of 14d organoid cultures derived using $Tdt^{+}$ AT2 cells from either *Abca3*$^{Cre}$*Rosa26*$^{Tdt}$ mice (labeled AT2) or *Sftpc*$^{I73T}$ mice (labeled activated AT2) and the four fibroblasts population isolated from WT or *Sftpc*$^{I73T}$ mice at 21d post tamoxifen induction. B-C) Bar graphs of differentially expressed (Differential expression calculated between fibrotic and alveolar fibroblasts FC>1.5 with adjusted p value <0.05) BMP signaling genes in fibroblasts subsets highlight a decrease in Bmp4 in inflammatory and fibrotic fibroblasts and an increase in BMP antagonists *Inhba*,*Grem1*, and *Fst* in these same populations. Additionally, we see a significant increase in *Tgfb1* and *Tgfb2* when performing direct ordinary one-way ANOVA analysis on inflammatory and fibrotic fibroblast normalized counts. Data is derived from population RNA-sequencing analyzed in Figure 4 (mean ± SEM; n=3 mice per group). D) Representative fluorescent microscopy images of 14d organoid cultures derived using $Tdt^{+}$ AT2 cells from *Abca3*$^{Cre}$*Rosa26*$^{Tdt}$ mice and the four fibroblasts population isolated from WT or *Sftpc*$^{I73T}$ mice at 21d post tamoxifen induction. All organoid cultures were supplemented with 5ng/ml mouse TGFβ1 from days 2-14 of the organoid culture. Scale bars:500μm. *p<0.05, **p<0.005, ***p<0.0005 by ordinary one-way ANOVA.

**Figure S6**

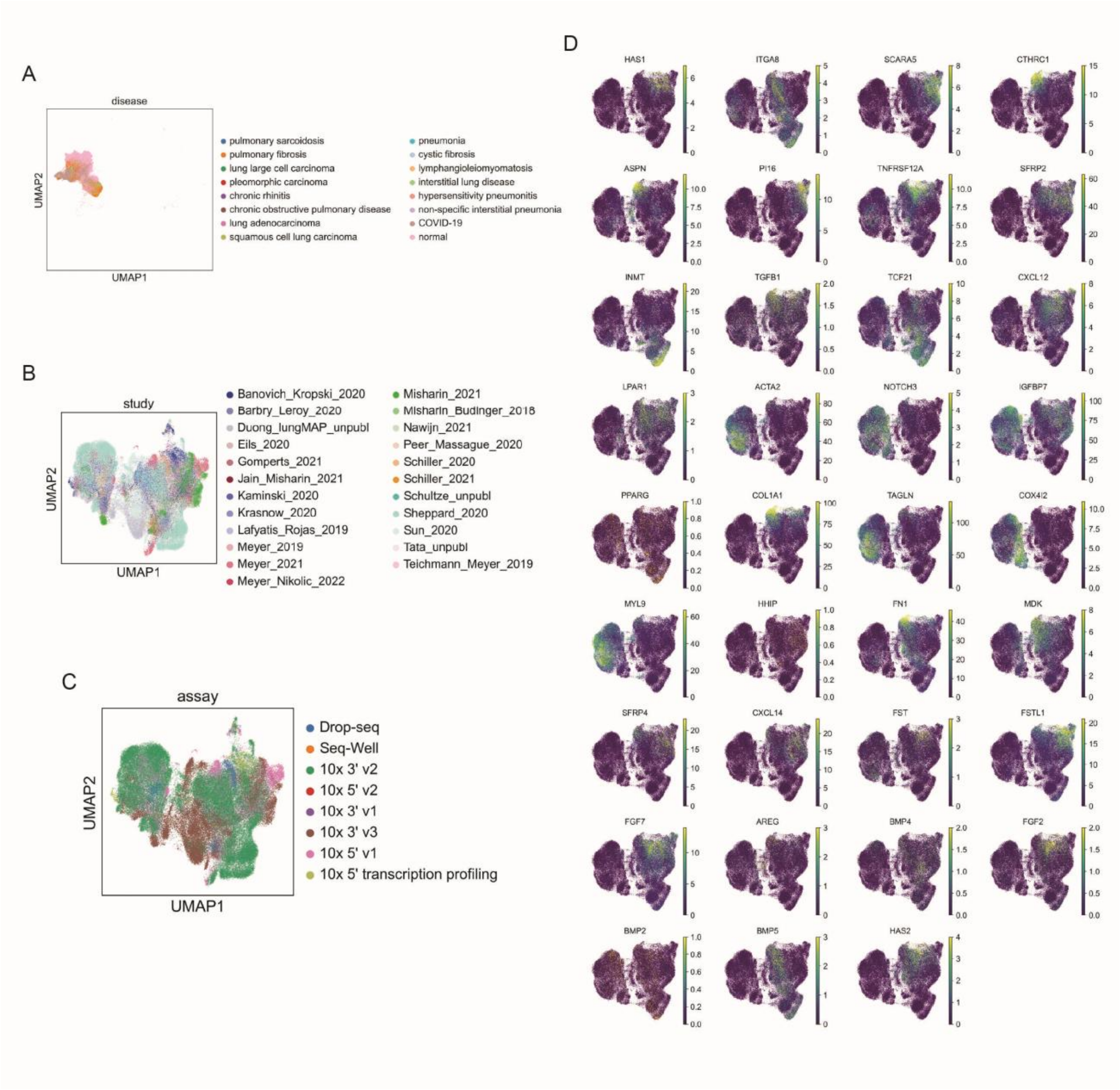
A
disease
UMAP2
UMAP1
pulmonary sarcoidosis
pulmonary fibrosis
lung large cell carcinoma
pleomorphic carcinoma
chronic rhinitis
chronic obstructive pulmonary disease
lung adenocarcinoma
squamous cell lung carcinoma
pneumonia
cystic fibrosis
lymphangioleiomyomatosis
interstitial lung disease
hypersensitivity pneumonitis
non-specific interstitial pneumonia
COVID-19
normal
B
study
Banovich_Kropski_2020
Barbry_Leroy_2020
Duong_lungMAP_unpubl
Eils_2020
Gomperts_2021
Jain_Misharin_2021
Kaminski_2020
Krasnow_2020
Lafyatis_Rojas_2019
Meyer_2019
Meyer_2021
Meyer_Nikolic_2022
Misharin_2021
Misharin_Budinger_2018
Nawijn_2021
Peer_Massague_2020
Schiller_2020
Schiller_2021
Schultze_unpubl
Sheppard_2020
Sun_2020
Tata_unpubl
Teichmann_Meyer_2019
C
assay
Drop-seq
Seq-Well
10x 3' v2
10x 5' v2
10x 3' v1
10x 3' v3
10x 5' v1
10x 5' transcription profiling
D
HAS1
ITGA8
SCARA5
CTHRC1
ASPN
PI16
TNFRSF12A
SFRP2
INMT
TGFB1
TCF21
CXCL12
LPAR1
ACTA2
NOTCH3
IGFBP7
PPARG
COL1A1
TAGLN
COX4I2
MYL9
HHIP
FN1
MDK
SFRP4
CXCL14
FST
FSTL1
FGF7
AREG
BMP4
FGF2
BMP2
BMP5
HAS2

**Figure S6. Analysis of stromal cells in the Human Lung Cell Atlas.** A) UMAP visualization of human mesenchymal cells from the Human Lung Cell Atlas and color-coded by disease status. In this plot the UMAP projection was not recalculated after removal of non-stromal cells. B) UMAP projection after removal of non-stromal cells and recalculation using scanpy.pp.neighbors and scanpy.tl.umap, color coded by studies included in this data set. C) UMAP projection color coded by sequencing technology D) UMAP representation of mesenchymal genes used to define all cell clusters in this mesenchymal data set.
